# Clockwork Orangutan: microRNAs, thermoregulatory tradeoffs, and models of brain size evolution

**DOI:** 10.1101/2024.05.21.595052

**Authors:** Thomas Sorger, Bastian Fromm

**Author notes:** Second corresponding author.

## Abstract

The Expensive Tissue Hypothesis was proposed to account for thermal homeostasis during the evolutionary expansion of brain size in the human lineage and abandoned following publication of a study that found no significant anticorrelations among the body mass-independent residuals of metabolically expensive organs in over 100 mammals, including 23 primates. Re-examination of the same dataset reveals a consistent tradeoff between the liver and brain proportions of a four-organ thermogenic core (kidney, heart, liver and brain), an inherent mechanism of thermoregulation that predates the emergence of permanent homeothermy. The ability of current models of brain size evolution to account for thermal homeostasis is limited by two common assumptions: that organ sizes evolve independently, and that the energy cost of the brain is proportional to the log ratio of brain mass with respect to body mass. Instead, arithmetic ratios provide direct experimental evidence for thermoregulatory constraints on brain size, as do organ cellular metabolic rates (*cMR*s). These are inferred without log transformation from the parallel adaptive increases in *MR*/kg and number of microRNA families (*mirFam*) that have accompanied major shifts in mammalian evolution. The *cMR* of the liver, the primary organ of gluconeogenesis, varies inversely with that of the brain, the primary consumer of glucose, a phylogenetic plasticity that appears to recapitulate the liver’s unique developmental plasticity. With *mirFam* as a proxy for energy reliability, a positive feedback model of relative brain size detected adaptation to a low-energy regime among the smallest primates and, among the largest primates, adaptation to the physiological limit on the rate of heat dissipation.

## INTRODUCTION

With the emergence of 24-hour homeothermy, Boreoeutheria crossed a threshold of energy reliability relative to earlier-branching placentals and marsupials, among which many orders exhibit some degree of heterothermy^1-2^. During the subsequent ecological diversification of Laurasiatheria and Euarchontoglires^3^, carnivores, rodents and primates exhibited consistent increases in relative brain mass^4^. The evolution of resting metabolic rate (*rMR*), in concert with that of thermal conductance (*C*_min_), allowed Boreoeutheria to occupy nearly every climatic zone regardless of size^5, 6^, albeit with a narrower range of tolerance to heat than to cold^7-9^, evidence that variation at the higher end of the ranges of body size and ambient temperature has been constrained by a physiological limit on the rate of heat dissipation (HDL)^10-12^.

Simulation of two million years of African and Eurasian climate and habitats supports the view that *H. sapiens* evolved from *H. heidelbergensis* in southern Africa during an unusually warm period^13^, consistent with the “Savanna hypothesis”^14-16^. The evolution of modern humans appears to have been conditioned by the HDL: among hunter-gatherers and contemporary indigenous populations, both resting and active *MR*s are actually elevated compared with those of non-human apes^17-18^, and these high rates of energy expenditure are associated with uniquely human adaptations that promote heat loss, such as hairlessness, bipedalism, and a 10-fold increase in the density of eccrine sweat glands relative to the densities found in chimpanzees and macaques^19^. The latter adaptation may have enabled early humans to run down game in long distance pursuits^20^, but may have also made them uniquely susceptible to dehydration^21^, a marathon-enabling tradeoff.

Another potential mechanism to address the thermoregulatory challenge of a larger brain, first proposed four decades ago, is a tradeoff in energy expenditure between the brain and another metabolically expensive tissue, such as kidney, heart, liver or intestine^22-23^. Attempts to detect a tradeoff predicted by this “expensive tissue hypothesis” were abandoned following publication of an influential report by Navarette et al (2011) based on 100 mammalian species, including 23 primates, that found no significant anticorrelation among the body mass-independent residuals of seven metabolically expensive organs^24^. However, re-examination of the same dataset reveals that, of six possible tradeoffs among the four most highly thermogenic organs (kidney, heart, liver and brain), the proportion of liver mass is strictly anticorrelated with that of the brain. Since this tradeoff is conserved across 104 placental and 4 marsupial species, it represents a previously unrecognized plasticity in the control of mammalian organ size^25^, an inherent mechanism of thermoregulation that predates the emergence of permanent homeothermy.

Current models of brain size variation have accounted for phylogenetic, but not developmental non-independence. Furthermore, we demonstrate that the relative energy consumption of the brain is proportional to its arithmetic ratio with respect to body size rather than the log ratio, as commonly assumed, yielding direct experimental evidence for a thermoregulatory constraint on brain size. We develop a model of organ rates of thermogenesis free of these assumptions based on the parallel adaptive shifts in resting metabolic rates (*rMR*s) and number of microRNA families that have characterized major stages in mammalian evolution: placentation, permanent homeothermy and the divergence of Catarrhini^12^.

MirGeneDB, the manually-curated library of microRNA genes^26-27^, has previously served as the reference microRNA database in a wide range of comparative^28-30^, phylogenetic^31-32^, developmental^33-34^ and evolutionary studies^35-36^. Carnivores and primates are characterized by higher body mass (*Mbdy*) and body temperatures (*T*_b_) and faster resting metabolic rates (*rMR*s) than marsupials or earlier-branching placentals (Atlantogenata)^1-2^, differences that coincide with an increased number of conserved microRNA families (*mirFam*)^26-27^ and density of microRNA target sites^12^. The expansion of brain size in the hominid lineage also depended on a significant increase in *rMR*^17-18^, and their clade rank order corresponds to the rank order of *mirFam*; conversely, the divergence of smaller-from larger-bodied clades has been repeatedly associated with decreased *mirFam*: Glires from primates, Strepsirrhini from Haplorhini and Callitrichinae from Platyrrhini^12, 26-27^. Thus, each increase in the range of *mirFam* has defined a new range of variation of *rMR* and *Mbdy* and, consequently, the potential range of brain size variation, reflecting two potential undelying mechanisms.

Cassidy et al (2019)^37^ investigated the nature of the relationship between *rMR* and *mirFam* during *Drosophila* development. While differentiation of the eye depended on microRNA activity in wild type *Drosophila*, development proceeded normally without any microRNAs when flies were engineered to have slower *MR*s, thereby doubling the time to eclosion. The authors reasoned that the normal unfolding of eye development must depend on the transient expression of transcription factors and the timely decay of their concentrations, implying that flies with wild type *MR*s and rates of protein synthesis are dependent on microRNA-mediated repression in order to maintain developmental synchrony^37^. By extension, the adaptive increases in *mirFam* that have accompanied mammalian shifts to faster *rMR*s may have also served to maintain synchrony among gene regulatory networks (GRNs). During mouse cranial neural crest differentiation, for example, specification of neuronal, as opposed to ectomesenchymal cells, depends on prolonging a transient period of chromatin accessibility in neuronal stem cells, an outcome of the repression of chromatin condensation by *Mir*-302^38^. Furthermore, since intrinsic protein expression noise increases with the rate of translation^39^, metabolic efficiency as well may have depended on matching faster *rMR*s with increases in global microRNA activity: protein synthesis drives 20% of cellular metabolic rate^40^ and global repression by microRNAs has been estimated to decrease the overall rate of translation by ∼ 25%^41-44^.

We postulated that the kidney, heart, liver and brain together serve the role of thermogenic “Core” predicted by a biophysical model of homeothermy^45^ and tested the following predictions:

i. differences in measured total energy expenditure among Hominoidae will reflect differences in the proportion of the brain mass and brain *rMR* in the Core;
ii. the arithmetic ratio of relative brain size will decrease with body size and evolutionary reductions in body size will be accompanied by increases in relative brain size, irrespective of phylogeny;
iii. Core organ cellular *MR*s (*cMR*s), optimally scaled with respect to body mass and *mirFam*^12^, will exhibit a joint rate of thermogenesis that scales as predicted by the biophysical model of endothermy (∼ *M*^0.50^)^45^.
iv. phylogenetic rates of evolution of liver and brain *cMR*s will reveal adaptive tradeoffs accompanying the divergence of clades between larger- and smaller-bodied mammals.

We demonstrate that the *mirFam*-based cellular *MR*s (*cMR*s) of the kidney, heart and brain vary directly with body mass, while liver *cMR* varies inversely with mass. Since the brain is the primary consumer of glucose and the liver is the primary organ of gluconeogenesis, these tradeoffs are consistent with a positive feedback model of brain size evolution governed by energy reliability, as has been previously suggested^4, 49^.

## RESULTS

### The nature of the 108-mammal dataset of organ mass and metabolic rate

For ethical and technical reasons there is a paucity of direct measurements of their oxygen consumption of the four primary thermogenic organs. Wang et al (2012)^50^ derived allometric models based on measurements of organ metabolic rates in the human, rat, rabbit, cat, and dog^50-52^, and applied them to the organ mass dataset of Navarette et al (2011)^24^, the same dataset that provided the basis for refuting the Expensive Tissue Hypothesis. Together with a term for residual thermogenesis, Wang et al (2012) fit these four organ *rMR*s to an additive model of whole-body *rMR*: *rMR* = 70 x *M*^0.75^. Henceforth, we refer to this dataset of measured organ sizes and predicted *rMR*s as the “Wang dataset”^50^.

### Tradeoffs among Core organs verify the expensive tissue hypothesis

Using the Wang dataset^50^, we obtained the organ/Core ratios of the measured mass of each thermogenic organ in 108 therian mammals (Figure 1-B). Of the six possible tradeoffs among four thermogenic organs, the most highly conserved was that between brain and liver, with a dampened tradeoff between heart and kidney. Brain/Core mass ratios ranged from 0.04 (pig) to 0.36 (human) to 0.70 (squirrel monkey), while Brain/Liver ratios ranged from 0.05 (opossum) to 0.76 (human) to 1.49 (squirrel monkey) (Figure 1-B). The variation in the proportions of the Core organs bore no apparent relationship to either body size or phylogenetic clade with the exception of the heart, whose relative proportion was low among the smallest rodents and high among the largest carnivores (Figure 1).

**Figure 1.**
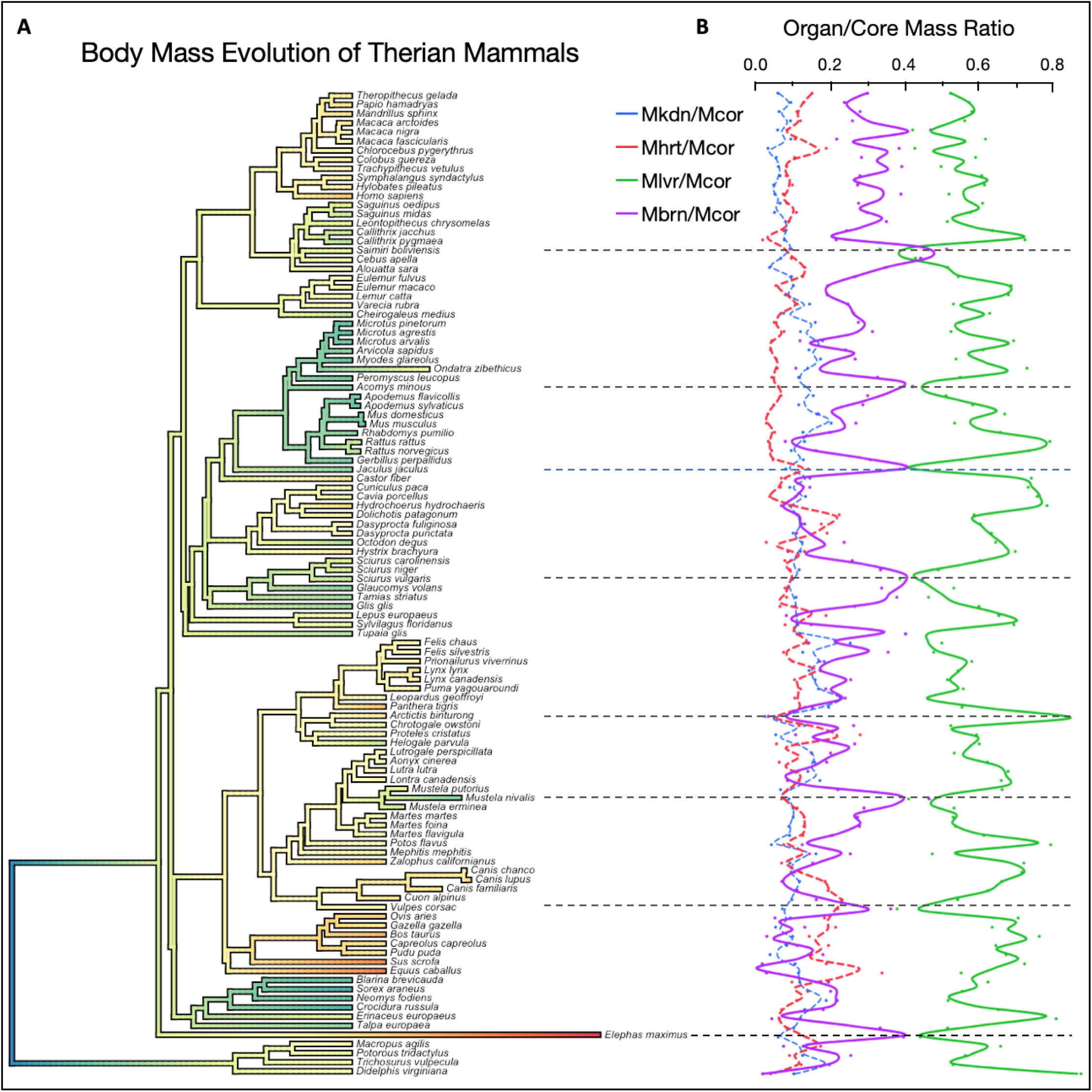
The proportion of brain mass in the thermogenic Core varies reciprocally with that of the liver. **A.** Variable rate phenogram of body size variation among Therian mammals. The color scale represents the five-fold range of log_2_ mass. Branch lengths were obtained by Bayesian inference with the expectation of a Brownian motion model of trait evolution^46-47^. Differences in root-tip branch length from the ultrametric tree^48^ represent either accelerated or decelerated rates of trait evolution. In the case of three outliers, branch lengths were arbitrarily shortened by 50% (*Ondatra zibethicus*, *Mustela nivalis*) or 80% (*Elephas maximas*). **B.** Mass proportion of each organ relative to the combined Core mass (*Mcor*). *Mkdn*, *Mhrt*, *Mlvr* represent the mass of the kidney, heart, liver and brain, respectively. The mass of each organ was divided by the combined mass of the four thermogenic organs in each species. Dashed lines indicate species with extreme ratios of brain mass/liver mass.

A tradeoff between liver and brain mass was also readily demonstrable in log-transformed data, provided that these were regressed on Core mass (Figure 2-B), rather than body mass (Figure 2-A): the Core mass-independent residuals for liver and brain were highly anticorrelated (*R*^2^ = 0.62, *P* = 2.32e-23) (Figure 2-B). The inability of the conventional analysis of brain size evolution, the phylogenetic regression of log *Mbrn* (brain mass) on log *Mbdy* (body mass), to detect such a highly conserved adaptation prompted us to investigate further the implications of two key assumptions of conventional phylogenetic comparative models.

a. While these methods correct for phylogenetic non-independence, they do not take into account developmental non-independence, *i.e.* the gene regulatory programs that ensure that mammalian organ sizes track body size during development^25^ (Supplementary Figure S1). For this reason, models of brain size standardized to body size will be blind to tradeoffs in the residual variation among the four thermogenic organs within the Core. Since whole-body *rMR* is the sum of organ *rMR*s, it is likewise not independent of body size, a shortcoming that is compounded when *rMRbdy* is treated as an independent variable in models of the relationship between log *Mbrn* and log *Mbdy*^4^.
b. An adaptation that releases a constraint on the rate of heat dissipation may allow a diverging clade to occupy a higher range of body mass, while the opposite adaptation might occur if the energy supply is no longer sufficient to maintain thermal homeostasis. For example, a recent model of stabilizing selection revealed two adaptive shifts in the evolution of primate brain size, corresponding to the divergence of larger-bodied Catarrhini from Haplorhini, and the subsequent divergence of smaller-bodied Callitrichinae from Platyrrhini (Figure 1)^54^. The resulting ranks of brain/body *arithmetic* ratios, Callitrichinae > other Platyrrhini > Cercopithecidae, were consistent with a disproportionate reduction in body mass (Figure 1, Inset), but contrary to the rank order of log ratios (Cercopithecidae > other Platyrrhini > Callitrichinae) (in both cases, *P* ≅ 0.0 for Callitrichinae *versus* Cercopithecidae: Dunnett’s Method with control). This interpretation is generalized by the arithmetic mass ratio of the 4-organ thermogenic Core (*Mcor*/*Mbdy*), which decreases strictly as a function of body size, consistent with a geometric constraint on the rate of heat dissipation (Figure 3).
c. The curvature of the relationship between log *Mbrn* and log *Mbdy* is *prima facie* evidence of adaptation to thermoregulatory constraints, but owing to the limitations enumerated above, sample sizes on the order of hundreds of species are required detect a departure from linearity that is statistically significant^4, 53^. A practical implication of this insensitivity is that unless the allometric model can resolve a 3% difference in slope (*M*^0.10^) in the range *M*^0.33^ - *M*^0.66^, a value that is likely to be within the range of measurement error, it will overlook a variation of 7% in the relative energy consumption of the brain, assuming that it is proportion to the ratio *Mbrn*/*Mbdy*.

**Figure 2.**
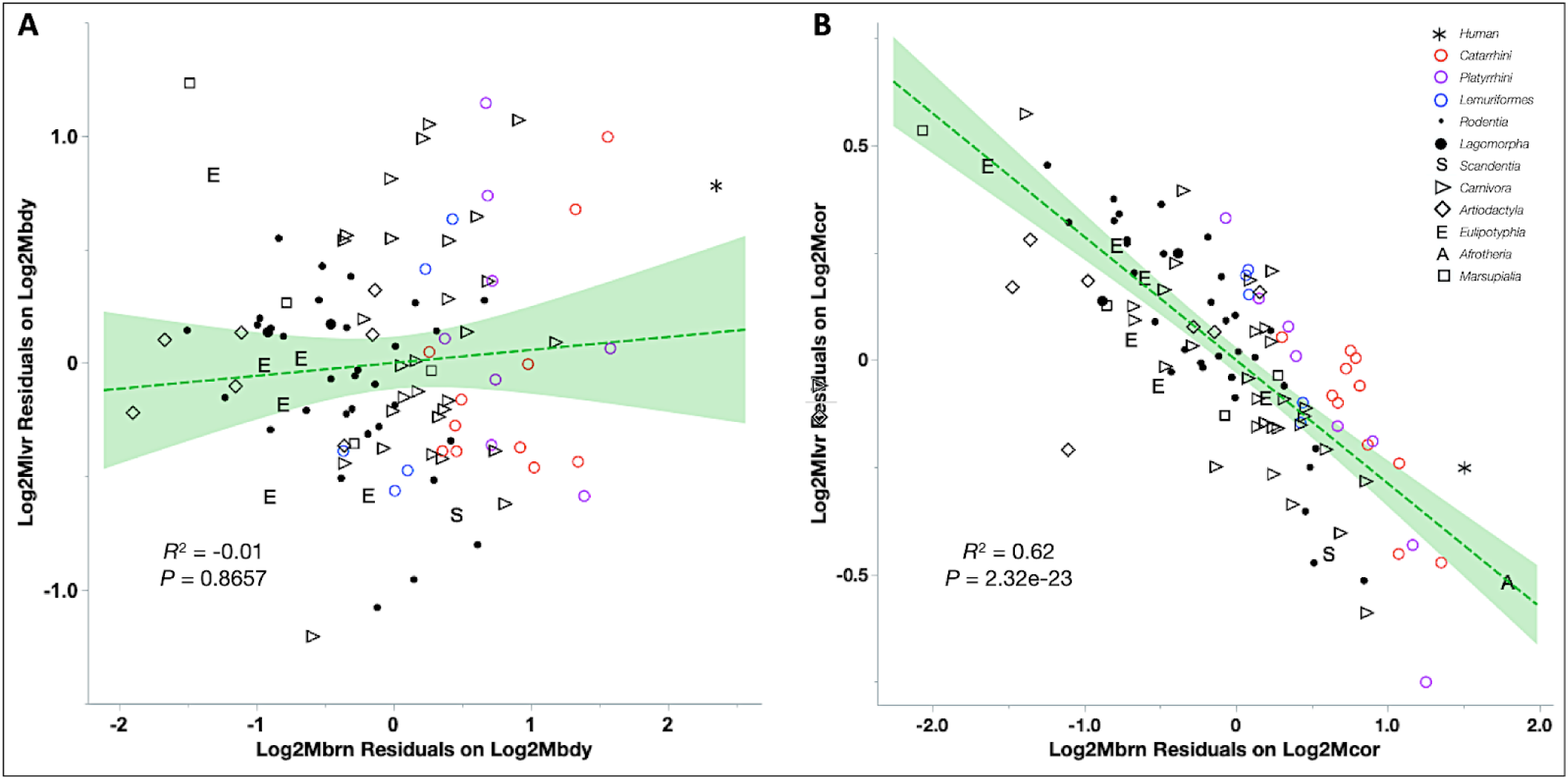
Variation of liver and brain mass relative to either whole body mass or the mass of the four-organ Core. **A.** Liver mass residuals with respect to body mass were plotted against brain mass residuals. **B.** Liver mass residuals with respect to Core mass were plotted against brain mass residuals. All 108 mammals in the dataset are shown, but the regressions represent only 88 carnivores, Glires and primates.

**Figure 3.**
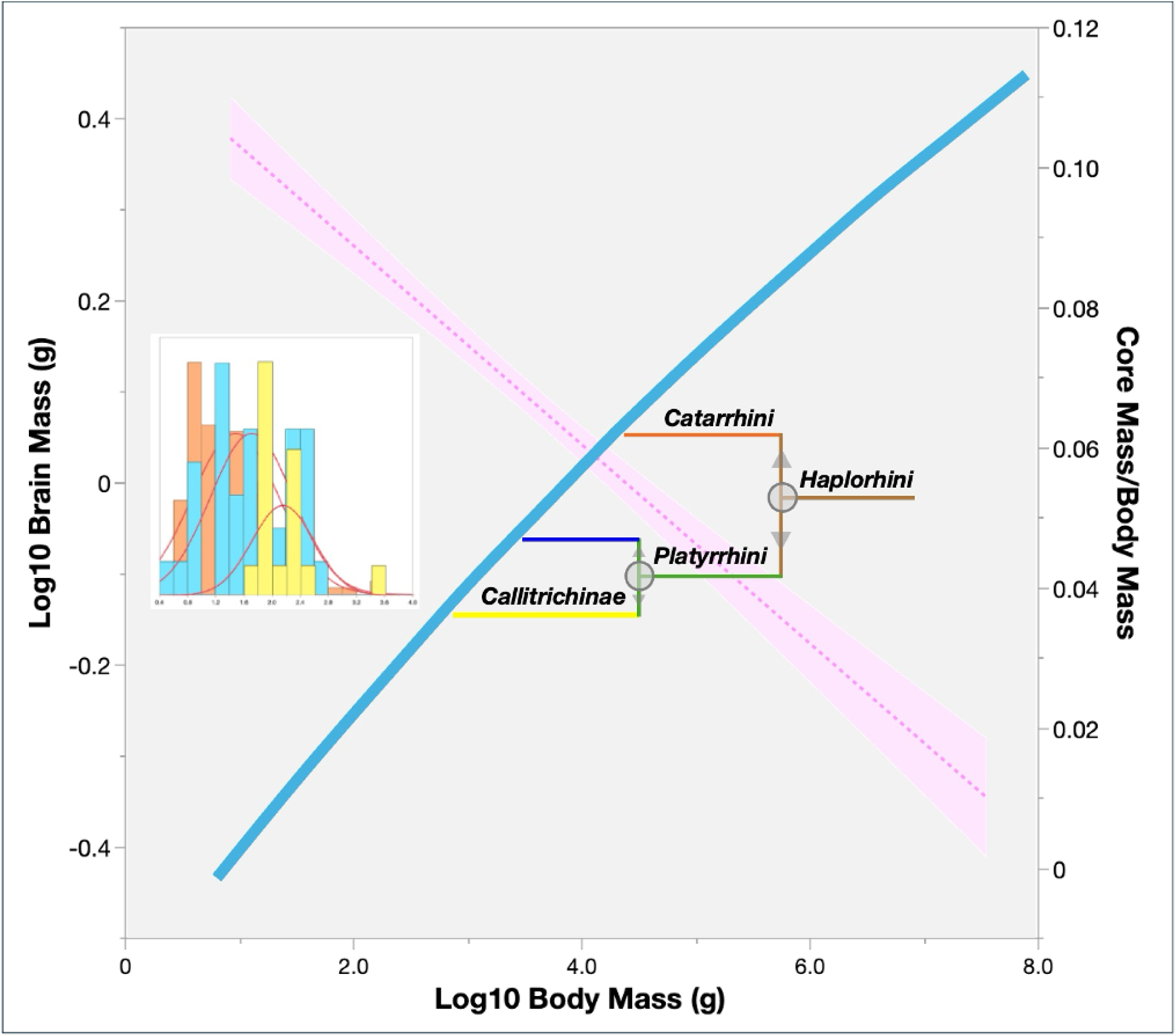
Evidence that clade shifts in the range of body mass represent thermoregulatory adaptations. The main curve describing brain size in 1500 mammals is derived from the quadratic model of Venditti et al (2024)^4^. The circles depict adaptive shifts in the evolution of primate brain size identified by Grabowski et al (2023)^54^. The pink dotted line and confidence band represent *Mcor*/*Mbdy*, the arithmetic ratio of the combined mass of the “Core” thermogenic organs to the body mass of 105 Carnivora and Euarchongotlires^50^ (*R*^2^ = 0.65, *P* = 2.23e-25). **Inset**: Brain/body mass ratios in Callitrichinae (yellow) > other Platyrrhini (blue) > Cercopithecidae (orange), consistent with a disproportionate reduction in body size (*P* = 0.0266 and 0.0, Dunnett’s Method with control). Data for Callitrichinae (N = 18), other Platyrrhini (N = 37), and Cercopithecoidea (N = 83): Grabowski et al (2023)^54^.

The combined mass of four thermogenic organs constitutes only 6% of mammalian body mass^50^, yet generates between 30 - 80% of the metabolic heat (55 ± 11.8%)^10^. The most significant limitation of current allometric models is that the evolution of brain size is treated in isolation from that of the other thermogenic organs. While developmental rules dictate that organ sizes track body size^25^ (Supplementary Figure S1), geometric constraints dictate that core thermogenesis decrease as body size increases (Figure 3).

### Differences among Hominoidae in the measured rates of energy expenditure reflect tradeoffs in the Core proportions of brain and liver mass and rMR

In order to assess the relevance of tradeoffs in the Core model to the evolution of brain size in the hominoid lineage, we compared the brain/Core and liver/Core ratios of mass and *rMR* with field measurements of the total energy expenditure (TEE) of orangutans and chimpanzees^17^. Using the double-labelled water method together with physical activity budgets, these authors found that the TEE of human males was 67% higher than that of *Pongo* and 25% higher than that of *Pan*, after correction for mass^17^. These differences mirrored a comparison of Core proportions between a set of three human samples *versus* two other hominoid species (*Symphalangus syndactylus* and *Hylobates concolor*) in the Wang dataset^50^: the brain/Core mass ratio was elevated 40% among humans, while the brain/Core *rMR* ratio was 68% higher (*P* = 0.0014 and 0.0004, respectively). These changes were offset by decreases of 16.4% in the liver/Core mass ratio and 29.6% in the liver/Core *rMR* ratio (*P* = 0.0033 and 0.0007, respectively; ANOVA with pooled t-test).

### The Inference of thermogenic organ cMRs

Previously, we developed an alternative to the log-based method for the estimation of specific *MR* (*sMR*) by fitting the distribution of *rMR*/*M*^exp^ to the distribution of microRNA families (*mirFam*), a proxy for the rate of energy turnover of a species^12^. Two advantages of this method are the greater resolution of the original (non-log transformed) values of *rMR* and the independence of *mirFam* from the developmental control of organ size. Thus, in contrast to the conventional estimation of a fixed *sMR* from the slope of log *rMRbdy* on log *Mbdy*, the distribution of organ *cMR*s is matched to the distribution of *mirFam* while controlling for the overall rate of thermogenesis, which is predicted to scale as *M*^0.50 45^.

Comparison of potential *cMR*s with allometric exponents across the range of 0.65 - 0.85 revealed opposite trends for liver, on the one hand (decreasing *P* value/increasing correlation), and for kidney, heart and brain, on the other (increasing *P* value/decreasing correlation), whether species were classified by range of *mirFam* or phylogenetic clade (Figure 4-A and B, respectively). Notably, the anticorrelation that emerges from this *mirFam*-based partitioning of organ rates of thermogenesis is independent of the anticorrelation revealed by the analysis of organ/Core mass ratios (Figures 1, 2). Assuming an operating range of *M*^0.70^ - *M*^0.80^, we assigned each organ a *cMR* corresponding to the lower limit of the curve of *P* values, hence:

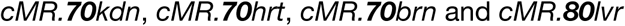

**Figure 4.**
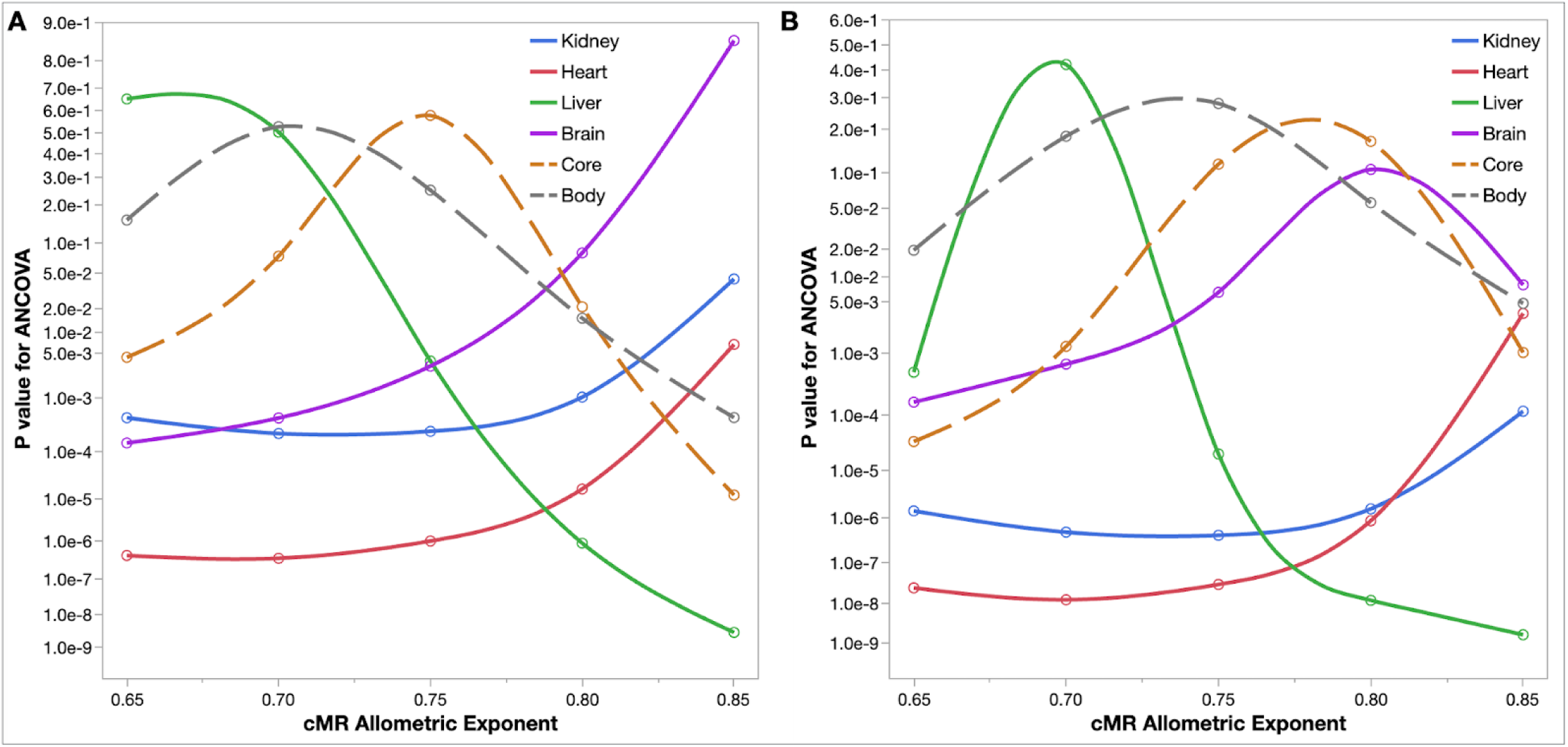
Inference of organ cellular *MR*s (*cMR*s): classification by *mirFam* range or phylogenetic clade. **A.** 50 Euarchontoglires were classified into bins according to natural divisions in the distribution of *mirFam*: 140-162 (Glires), 164-182 (Glires, Lemuriformes, Platyrrhini), and 194-210 (Catarrhini). *P* values represent regressions on Bin x √Mbdy (ANCOVA). Note that the interval *M*^0.70^ - *M*^0.80^ defines a zone of minimal correlation for the Core *cMR,* centered at *M*^0.75^, which is also the midpoint of the dynamic ranges of liver and brain *cMR*. **B.** The same analysis for species classified instead by phylogenetic clade (Glires, Lemuriformes, Platyrrhini, Catarrhini).

Whole body *cMR* primarily tracked liver *cMR*, while Core *cMR* exhibited a symmetric profile with a zone of minimal correlation centered at *M*^0.75^ (Figure 4-A). Since *M*^0.75^ was also the optimal allometric scaling in a model of stabilizing selection of mammalian *cMR* with respect to *mirFam*^12^, this zone of minimal constraint on the rate of Core thermogenesis may be interpreted as an evolutionary adaptation for thermal homeostasis. The same trends were observed when species were classified instead by phylogenetic clade, but in this case the curves for Core and whole body *cMR* were both shifted toward higher allometric exponents (Figure 4-B).

In order to rule out the possibility that the opposite patterns of correlation of liver and brain *cMR* are unique to the sample of variation represented by the Wang dataset, we applied the same allometric models to a completely independent set of *rMR*s measured in 30 primates^10^. Core *rMR* was inferred from its relationship to body *rMR* and then liver and brain *rMR* were inferred from Core *rMR* (Supplementary Table S3).

As in the case of organ/Core mass ratios, the *rMR*/Core ratios of liver and brain were anticorrelated in all but the largest primates (Figure 5-A). Furthermore, liver and brain *cMR*s exhibited opposite correlation patterns with respect *mirFam* and √*M*, and Core *cMR* exhibited minimal correlation when scaled to *M*^0.75^ (Figure 5-B). Thus, the observed anticorrelations between liver and brain are a consistent outcome of the Wang allometric models, regardless of species sampled or whether the source data represent organ mass or whole-body *rMR*.

**Figure 5.**
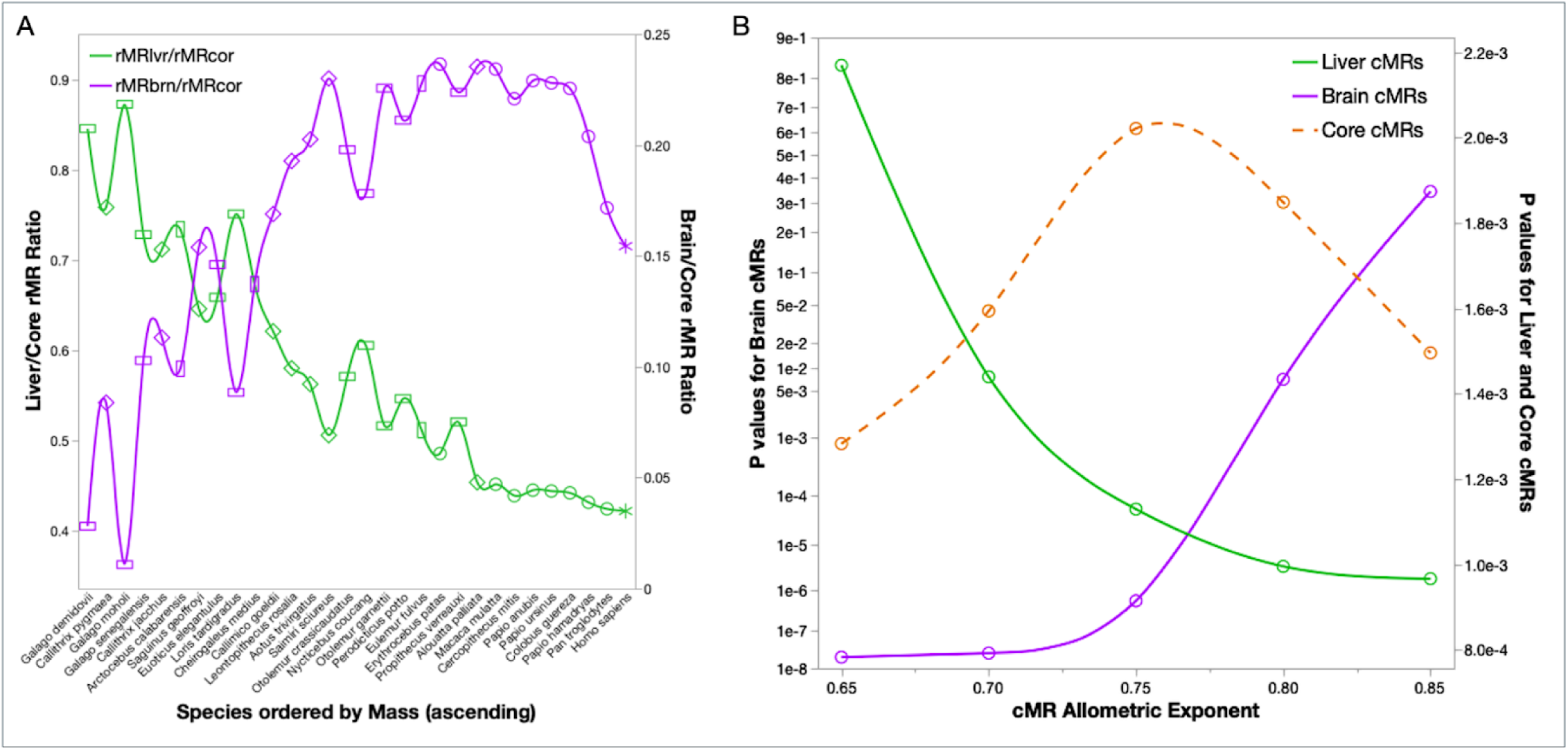
Core and organ *rMR*s can be inferred from measured primate *rMR*s. Based on the allometric models for 22 primates, excluding the human^13^, the following chain of inference was applied to the measured *rMR*s of 30 primates^10^ (Supplementary Table S3): organ mass ← organ *rMR* ← Core *rMR* ← Body *rMR*. **A.** Anticorrelation between organ/Core *rMR* ratios from independently derived liver and brain *rMR*s. **B.** Liver and brain *cMR*s exhibit opposite patterns of correlation with *mirFam* x √*Mbdy*. Core *cMR*s exhibit minimal correlation with *mirFam* x √*Mbdy* in the range *M*^0.70^ - *M*^0.80^ (compare with Figure 3).

### Organ cMRs, body mass, and thermal conductance

Plotting organ *cMR*s as a function of body size again revealed an inverse relationship between *cMR.80lvr* and the *cMR.70*s of the other three organs: while these increased directly with size, *cMR.80lvr* was strictly anticorrelated with body mass (Figure 6-A). A finer distinction was also evident: while the slope of *cMR.80lvr* was unchanged between smaller-bodied Glires and larger-bodied primates, the slopes for kidney, heart and brain *cMR.70* were significantly higher in the larger-bodied clade (Figure 6-A).

**Figure 6.**
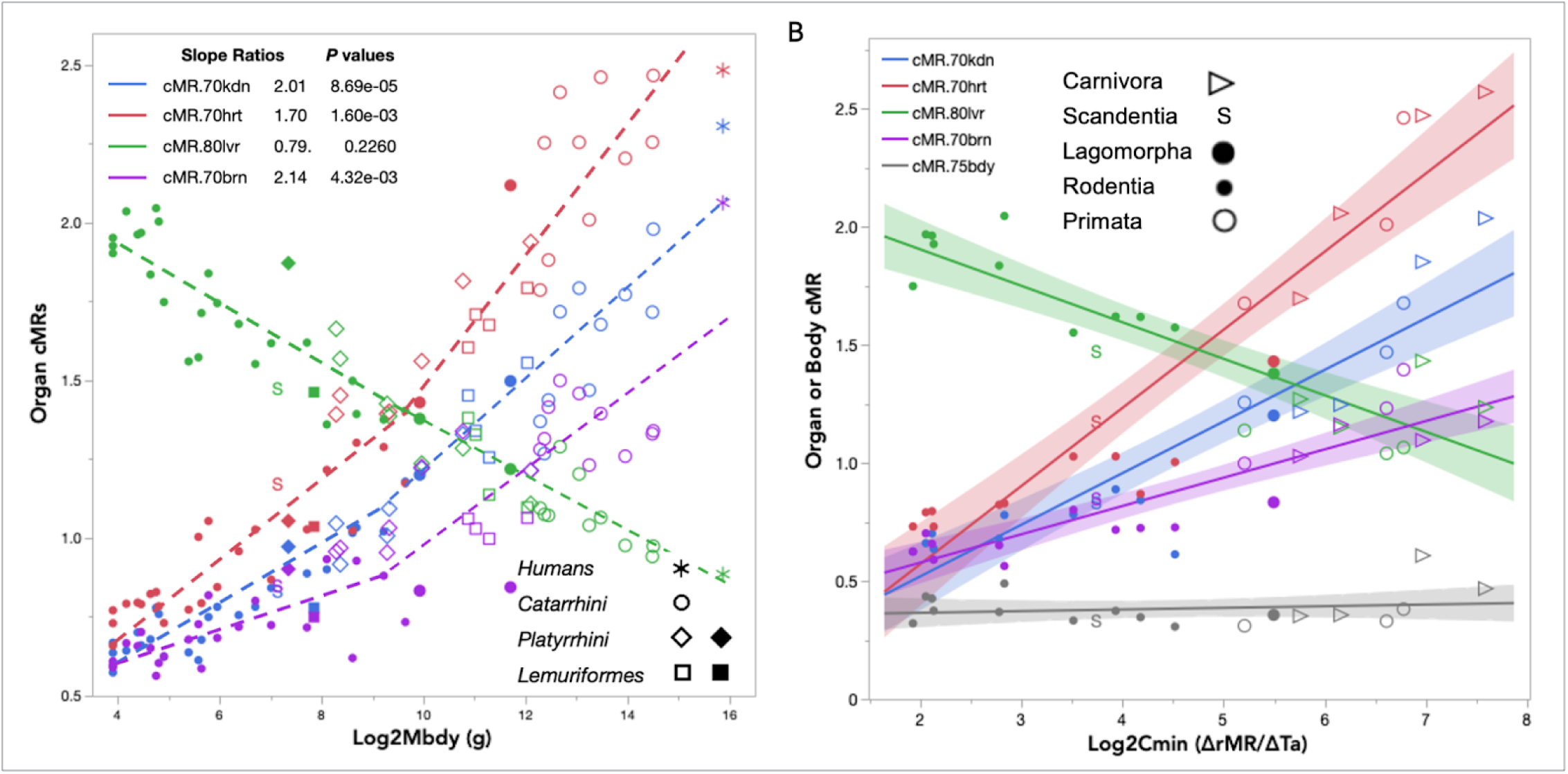
Core organ *cMR*s as a function of body size or thermal conductance (*C_min_*). **A.** Slope ratios between primates and Glires were compared by ANCOVA (figure legend) (N = 51). **B.** Organ rates of thermogenesis as a function of thermal conductance among Boreoeutheria (*N* = 20). *T*_a_ = ambient temperature. Shaded bands: confidence intervals (CI).

Not surprisingly, the same ranking of slopes (heart > kidney > brain) and anticorrelation with liver *cMR.80* was obtained when organ *cMR*s were regressed on the thermal conductance (log *C*_min_) of 20 Boreoeutheria (Figure 6-B). *C*_min_ is an extensive property that depends primarily on total body surface area and secondarily, on the properties of the insulating layer^45, 55^. When Boreoeutheria were ranked according the whole-body rate of heat generation (*cMR.75*), *mirFam* covaried with body size, core thermogenesis (*cMR.53cor*) and thermal conductance (log *C*_min_), independent of phylogeny (Supplementary Figure S2-A). As in the case of body size, ranking Boreoeutheria by *C*_min_ revealed tradeoffs between liver *cMR.80*, on the one hand, and both brain and heart *cMR.70*, on the other (Supplementary Figure S2-B), consistent with the coordinated evolution of body temperature, metabolic rate and thermal conductance^5-6, 56^.

### Thermodynamic and phylogenetic relevance of Core organ cMRs

We tested two predictions, the first stemming from the allometric model that was the basis of the *rMR*s in our dataset^50^, and the second from the thermoregulatory core model of Porter and Kearney (2009)^45^:

i. The joint ***sMR*** of the thermogenic organs should scale as *M*^0.75^, since the individual organ *rMR*s were inferred from an additive model assuming that whole body *sMR* = 70 x *M*^0.75^, and we ignored the residual *rMR*, which averaged only 4.5% of that of the liver in mammals < 75 kg^50^;
ii. The joint ***cMR*** of the four most highly thermogenic organs, although delocalized, should scale approximately as the geometrically centered, thermogenic core of the biophysical model *i.e.* ∼ *M*^0.50^.

Applying partial least squares regression^58^, we generated additive models in order to predict the optimal scaling of a joint Core *cMR* composed of either log-based *sMR*s:

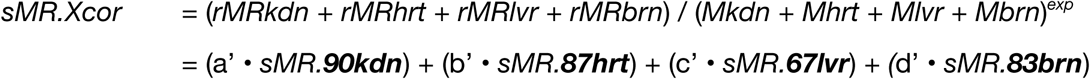

*or mirFam*-based *cMR*s:

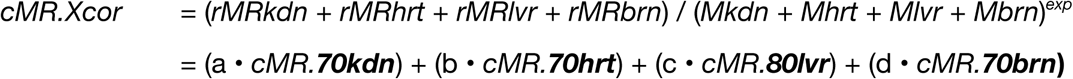

The log-based joint *sMR* exhibited a single sharp peak at *M*^0.75^, corresponding exactly to the whole-body *sMR* of both the biophysical model^45^ and the Wang allometric model from which the organ *rMR*s were derived^50^. The composite *cMR* model exhibited two peaks, one on either side of an inversion of Core *cMR* slopes with respect to √M: a broad peak centered at *M*^0.53^, similar to that predicted by the biophysical model of core thermogenesis^45^, and a second peak between *M*^0.80^ - *M*^0.85^ (Figure 7-A). Importantly, the *cMR* and *sMR* models exhibited critical differences in *VIP* statistics (*Variable Importance of Projection*), a measure of the model dependence on each variable^57^, analogous to the loadings of principal components. The *cMR* models depended on all four organ *cMR*s across the entire range, with the exception of a zone of minimal correlation (*M*^0.75^ - *M*^0.80^), where they depended only on that of liver. In contrast, the *sMR* model depended only liver, heart and brain (*M*^055.^ - *M*^0.65^) or only liver and kidney (*M*^0.70^ - *M*^0.90^) (Figure 7-A, bottom).

**Figure 7.**
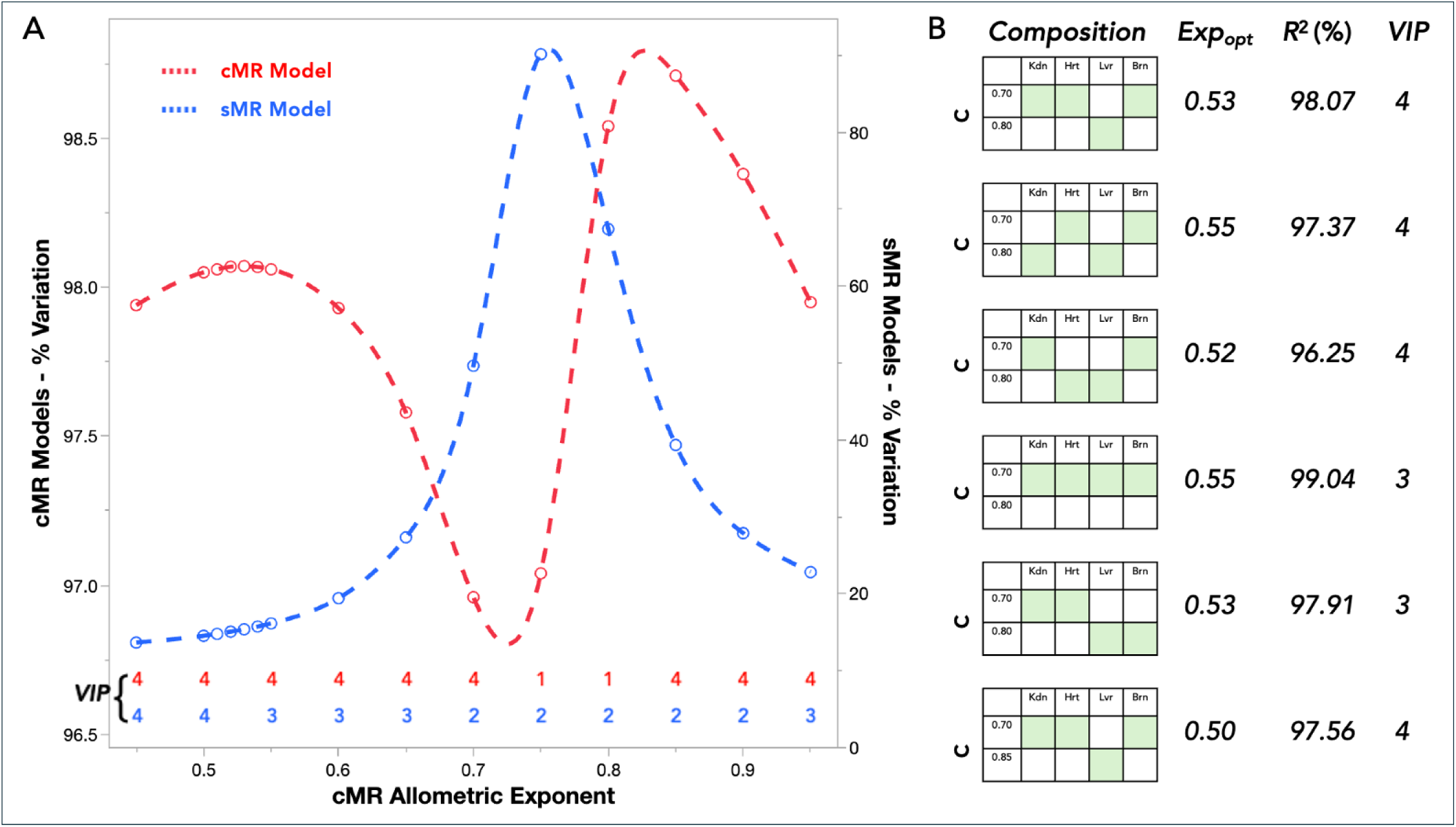
Scaling of the Core *cMR* and body *sMR* by linear combinations of either the *mirFam*-based *cMR*s or the log-based *sMR*s of the four thermogenic organs (*N* = 22 primates). **A.** % variation (*R*^2^) of linear models of Core *cMR* or body *sMR* composed of the four organ *cMR*s or *sMR*s. *VIP* values (*variable importance of projection*) indicates the number of organs upon which each model depends^57^. **B.** Comparison of the joint *cMR* of combinations of *cMR.70* - *cMR.85* among the four thermogenic organs. “*c*” is the organ *cMR* allometric exponent. *Exp*_opt_ is the optimal allometric exponent of the additive model.

We applied an additional test of the relevance of these estimated organ *cMR*s by comparing the joint *cMR*s obtained by permuting combinations of estimated organ *cMR*s, yielding four models with *VIP* = 4 (Figure 7-B). Of these, we found two combinations that were able to resolve Platyrrhini from Lemuriformes in an unsupervised classification of major primate clades^59^: either *cMR.80* or *cMR.85* for liver, together with *cMR.70* for heart, kidney and brain. When liver *cMR* was varied between *cMR.75* - *cMR.85*, the original set of *mirFam*-based *cMR*s yielded the lowest value for the Bayes information criterion (BIC) (Supplementary Figure S3), hence the following optimal model of the Core *cMR*:

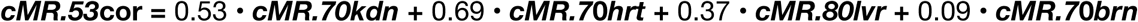

In the context of the evolution of thermal homeostasis, the coefficient of each organ *cMR* in this model indicates the degree to which it has been constrained by natural selection, *i.e.* the smaller the coefficient, the greater the permitted variation of the organ *cMR* since clade divergence.

The original biophysical model for thermoregulation assumed that metabolic heat emanates from the geometric center of an ellipsoid endotherm. Depending on the wind condition or the presence of fur, the authors derived a single allometric exponent between 0.46 - 0.58 in order to scale the rate of core thermogenesis across the entire range of mammalian body size^45^. Since our Core model is delocalized among four thermogenic organs and makes no geometric assumptions, we expected lower allometric exponents and higher rates of thermogenesis (*rMR*/*M*^c^) in smaller mammals with higher ratios of surface area/volume.

The optimal allometric slopes of *cMR* models developed in Carnivora, Primata and Glires were *M*^0.60^, *M*^0.53^ and *M*^0.40^, respectively (Figure 8). The corresponding distributions of *Mcor*/*Mbdy* support the notion of geometric constraints: Glires occupy the upper end of the range, suggesting an adaptation to maintain body temperature, while primates and carnivores occupy the lower end, consistent with a limit on the rate of heat dissipation (Supplementary Figure S4-A)^11^.

**Figure 8.**
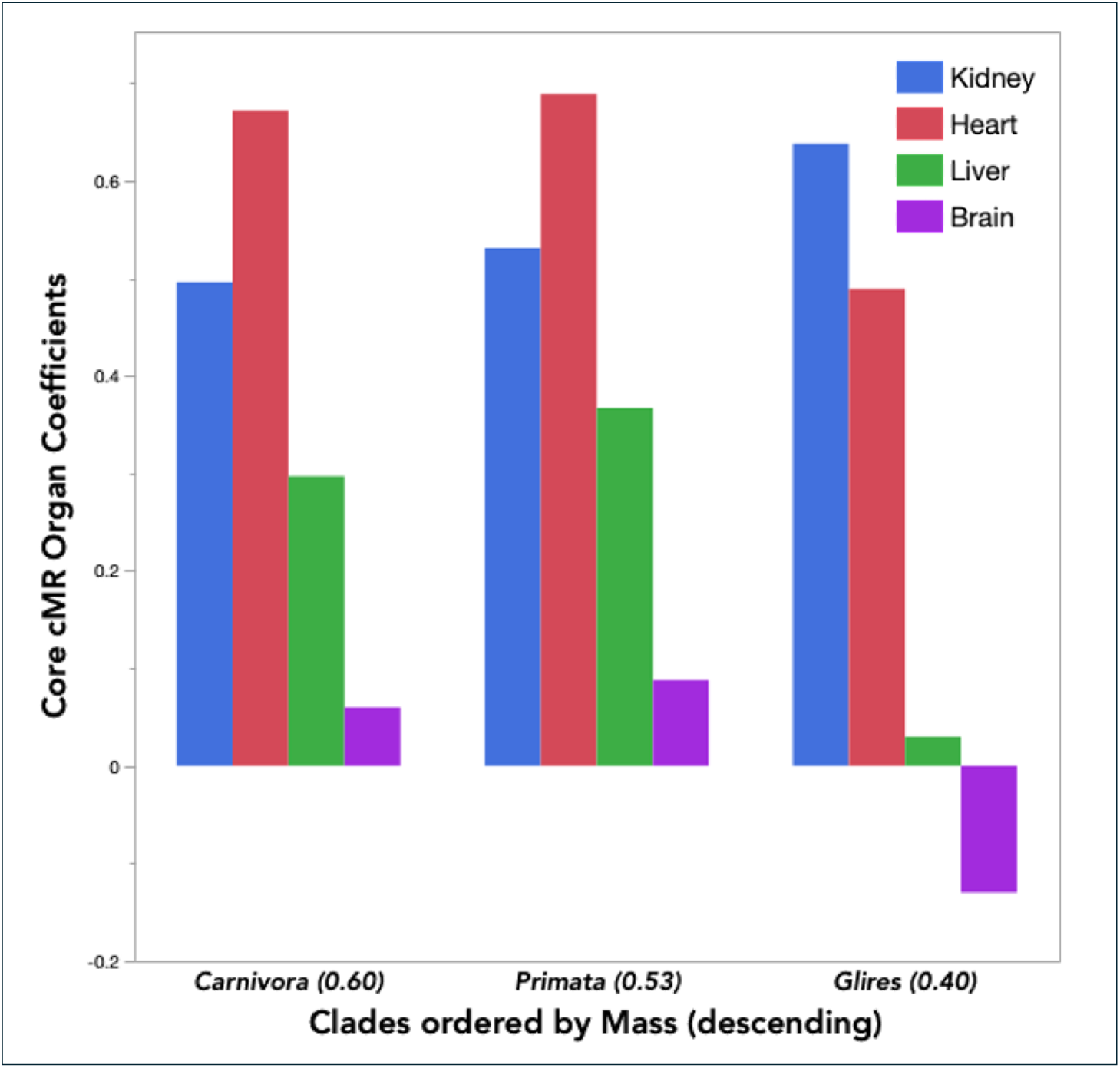
Organ coefficients of additive models of Core thermogenesis in three major clades of Boreoeutheria. Values in parentheses next to each clade indicate the optimal allometric exponent of each Core *cMR* model.

The coefficients in the Glires Core model present a striking contrast with those of carnivores and primates: a reversal in sign of the coefficient for brain *cMR* in Glires, and a reversal of the relative magnitudes of the coefficients for liver and brain (Figure 8). The Glires model thus penalizes the *cMR* of the brain, suggesting adaptation to a disproportionate reduction of body size relative to brain size, leaving the arithmetic mass ratio, *Mbrn*/*Mbdy,* higher in Glires than in primates or carnivora (*P* < 0.001, analysis of means) and contrary to the rank order of the allometric slopes for log *Mbrn versus* log *Mbdy* (primates > Glires) (Supplementary Table S4).

### Tradeoffs and adaptations in the rates of evolution of organ cMRs

The preceding evidence suggests that Boreoeutheria have been able to maintain thermal homeostasis while diversifying body mass by fine-tuning the relative sizes of the four Core organs in relation to that of the liver. We applied Bayesian inference in order to investigate the historical timing of the corresponding tradeoffs.

In agreement with the Core *cMR* models, we found opposite directions of evolution between the organ *cMR.70*s that vary directly body size (kidney, heart and brain) and *cMR.80lvr*, which varies inversely with body mass, both within and between major clades (Figure 9). These observations are supported by a computational model that found that following an overnight fast, rates of liver gluconeogenesis, lipolysis and proteolysis were all elevated ∼ 10-fold in mice, relative to the corresponding rates in humans^60^.

**Figure 9.**
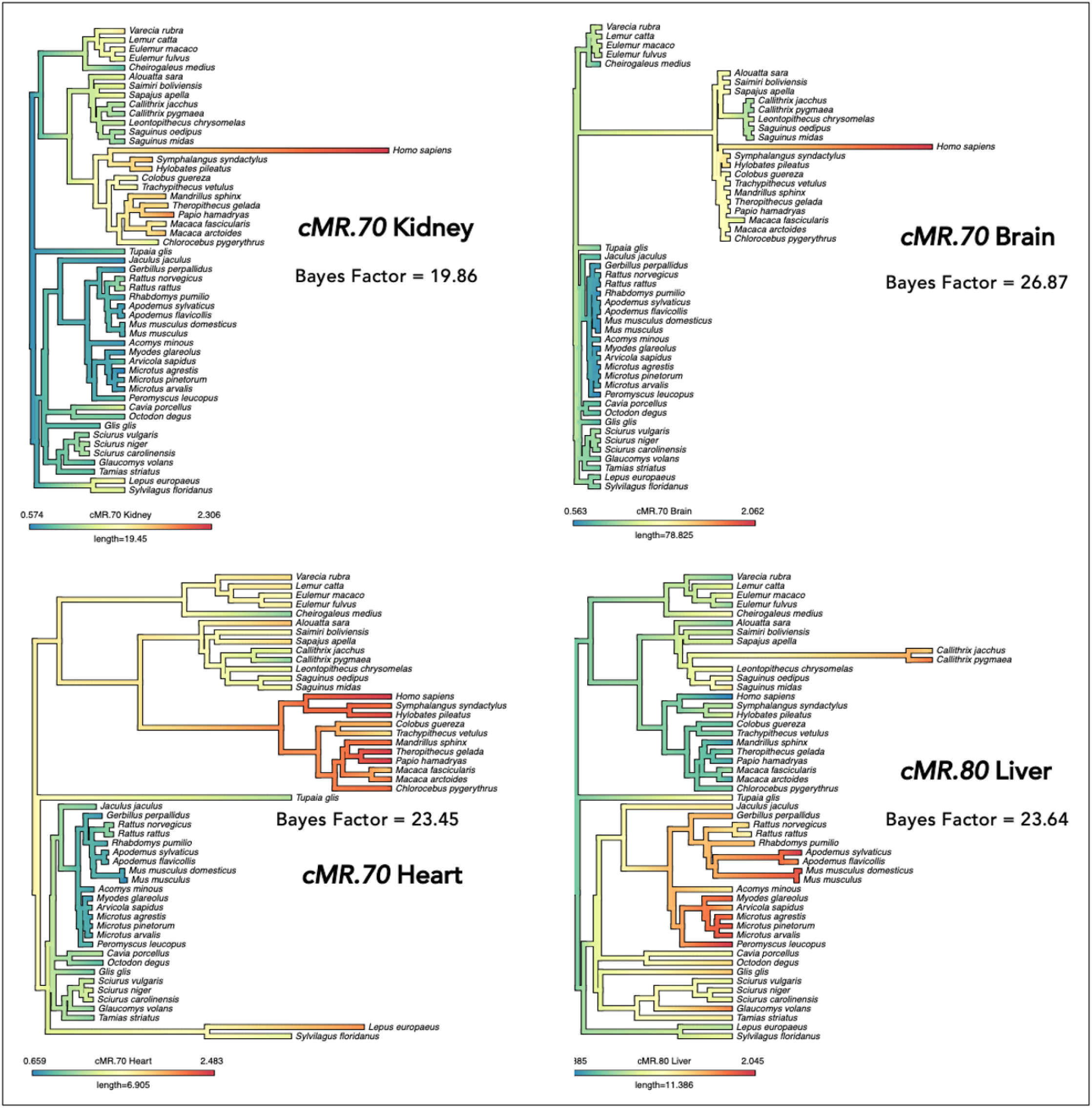
Variable rate phenograms of Liver *cMR* (*cMR.80*) and the *cMRs of* Kidney, Heart and Brain (*cMR.70*). Bayes factors were computed from the difference in log likelihood (*Lh*) of the 1% most likely trees and the trees with the smallest non-zero sum of changes to branch length^12^, *i.e.* those trees most similar to the reference time tree^48^.

Among primates, increased liver *cMR* accompanied the decrease in size of two of the smallest Platyrrhini, one of which (*Callithrix pygmaea*) may be heterothermic^61^. A clade of five small (< 1.0 kg) Platyrrhini exhibited an accelerated *decrease* in brain *cMR.70* paired with elevated levels of *cMR.80lvr*, consistent with adaptation to a low energy regime and greater reliance on gluconeogenesis. Among Glires as well, those species that underwent the most severe reductions in size exhibited the greatest increases in liver *cMR.80*; conversely, within Glires and Haplorhini, the large-bodied clades (Lagomorpha and Catarrhini) exhibited lower liver *cMR*s (Figure 8). Brain *cMR.70* increased in intensity and its rate of evolution accelerated with the divergence of Haplorhini, and then again in humans, in contrast to its stasis or decrease among Glires, Lemuriformes and smaller Platyrrhini (Figure 9). The relative levels of heart *cMR.70* generally tracked body size, with accelerated rates of heart evolution in the clades of the largest Glires (Lagomorpha) and primates (Catarrhini).

The divergence of humans was marked by the accelerated rate of evolution of the *cMR.70* of the kidney in addition to that of the brain (Figure 9), an observation that is noteworthy for two reasons:

a. The 30% higher level of kidney *cMR* found in humans relative to other hominoid apes is not apparent when kidney *MR* is scaled according to the log-based slope (*M*^0.90^): the corresponding value for the kidney *sMR* is 10% *lower* in humans than the average value among apes.
b. The adaptive increase in *cMR.70kdn* is of particular interest since the ability to thermoregulate is highly dependent on water balance, as illustrated in a study of baboons whose fossils appear in the same arid horizons as the fossils of *Australopithecus*^62^ and *Homo*^15^. When deprived of water and exposed to elevated ambient temperatures, free-ranging baboons developed a progressive hyperthermia (increasing core temperature) that was rapidly reversed upon rehydration^63^.

## DISCUSSION

### Direct experimental evidence for a thermoregulatory constraint on brain size

It now appears well established that a single rule^4^ governs the relationship between brain mass and body mass in mammals, irrespective of phylogeny, rather than multiple, clade-specific rules^64^. This paradigm comports well with the notion of a single developmental program in mammals to scale organ sizes to body size^25^. However, current phylogenetic comparative models have overlooked a thermoregulatory tradeoff embedded in this developmental program, an oversight the present study aims to correct. In order for the variation of a trait to be shaped by thermoregulatory constraints, it must reflect the relative energy cost *in vivo*, *i.e.* the arithmetic proportion of mass or heat production, rather than the log-based allometric slope: while log-based allometric slopes are positive, consistent with the developmental programming of organ sizes, Core/body mass ratios decrease strictly as a function of size, consistent with geometric constraints.

Venditti et al (2025) recently demonstrated that, irrespective of phylogeny, a quadratic function best fits the relationship between log brain mass and log body mass in 1,500 mammals, evidence for a universal constraint on evolution of brain size at the upper end of any range of body mass^4^. Hypothetically, this constraint could be either anatomic *i.e.* the diameter of the skull in relation to that of the birth canal, or physiological, such as a limit on the rate of heat dissipation^11^.

In order to test whether this constraint is related to the overall rate of thermogenesis, we inferred *rMR*s for the body mass data in a dataset of endocranial volumes (*ECV*)^54^ based on a separate set of *rMR*s measured in the same primate species^10^. Bayesian regressions were then carried out to compare the dependence of log *ECV* on log *Mbdy* and log *rMR* for 15 species in each of three, overlapping ranges of relative brain size (*ECV*/*Mbdy*). As shown in Table 1, the range of *ECV/Mbdy* was inversely related to the range of body mass. In each case, the model for the dependence of *ECV* on body mass was improved by the inclusion of the *rMR* term. For the highest range of body mass and lowest brain/body ratios, the regression coefficients for the *rMR* term were consistently (100%) negative, corresponding to 12.2% of the value of the *Mbdy* term. For the smallest-bodied species with the highest brain/body ratios, the regression coefficients for the *rMR* term were consistently (100%) positive, equivalent to 10.5% of the value of the *Mbdy* term. For the middle range of body size and relative brain size, the regression coefficient for the *rMR* term fluctuated around 0, as was the case for the entire dataset of 25 species. Thus, after correction for phylogenetic covariance, relative brain size was inversely related to metabolic rate in the 15 largest species, *i.e.* those most likely to have adapted to a heat dissipation limit. On the other hand, the *rMR* term was a positive function of relative brain size in the 15 smallest species, consistent with the greater relative energy demand of the brain, and for whom their surface area and rate of heat loss may have required adaptation in order to maintain body temperature.

**Table 1.** Sliding window analysis of the interaction between relative brain size and whole-body metabolic rate. The measured *rMR* data for 25 primate species from Clarke et al (2010)^10^ were matched with endocranial volumes (*ECV*) from Grabowski et al (2023)^54^. An exponential model of *rMR* ∼ *Mbdy* was applied to the body mass values of the *ECV* dataset. Bayesian phylogenetic regression was applied to three overlapping sets of 15 species ranked by *ECV*/*Mbdy* ratio in order to test the dependence of Log_2_*ECV* on Log_2_*Mbdy* and Log_2_*rMR*. The first bin in each distribution represents five species. All statistics represent the median value of the 100 regressions with the greatest likelihood (top 10%). The primate species included 6 Catarrhini, 5 Platyrrhini, 4 Lemuriformes and 10 Lorisiformes (Supplementary Table S6).

| Sample | Range Mbdy (g) | Range Ratio (%) | Likelihood | Model | $\beta_1$ | $\beta_2$ | $R^2$ |
| --- | --- | --- | --- | --- | --- | --- | --- |
| Low ECV/Mbdy (N = 15) | 140 – 22,300 | 0.80 – 1.86 | -5.31 | $\text{Log}_2\text{ECV} = \beta_1 \cdot \text{Log}_2\text{Mbdy}$ | $0.66 \pm 0.077$ | | 0.85 |
| | | | -3.08 | $\text{Log}_2\text{ECV} = \beta_1 \cdot \text{Log}_2\text{Mbdy} + \beta_2 \cdot \text{Log}_2\text{MR}$ | $0.58 \pm 0.079$ | $-0.07 \pm 0.060$ | 0.84 |
| Middle ECV/Mbdy (N = 15) | 140 – 14,150 | 1.02 – 2.26 | -6.22 | $\text{Log}_2\text{ECV} = \beta_1 \cdot \text{Log}_2\text{Mbdy}$ | $0.69 \pm 0.072$ | | 0.88 |
| | | | -5.95 | $\text{Log}_2\text{ECV} = \beta_1 \cdot \text{Log}_2\text{Mbdy} + \beta_2 \cdot \text{Log}_2\text{MR}$ | $0.70 \pm 0.074$ | $-0.04 \pm 0.057$ | 0.88 |
| High ECV/Mbdy (N = 15) | 75 – 989 | 1.49 – 3.59 | -3.19 | $\text{Log}_2\text{ECV} = \beta_1 \cdot \text{Log}_2\text{Mbdy}$ | $0.63 \pm 0.067$ | | 0.88 |
| | | | -1.88 | $\text{Log}_2\text{ECV} = \beta_1 \cdot \text{Log}_2\text{Mbdy} + \beta_2 \cdot \text{Log}_2\text{MR}$ | $0.67 \pm 0.069$ | $0.07 \pm 0.047$ | 0.90 |

### The developmental and phylogenetic plasticity of the liver

The brain and liver, each accounting for 20% of *bMR* in humans, exhibited highly conserved tradeoffs in mass and *cMR*, a plasticity that appears to be an inherent property of the mammalian evolutionary toolkit, not exclusive to homeotherms. While the negative slope for the dependence of liver *cMR* on body mass remained constant across clades, the positive slopes for kidney, heart and brain *cMR*s were significantly higher in primates than among Glires, suggesting that their combined arithmetic proportion has been adjusted relative to that of the liver in a clade-specific manner.

An inverse relationship between gluconeogenesis and the rate of body mass growth in humans implicates the liver as the pivotal partner in an ontogenetic tradeoff: brain glucose utilization peaks around age five, early in the extended period between weaning and sexual maturity, which corresponds to the period of slowest growth in body mass^65^. Based on the turnover of isotopically-labeled glucose, liver glucose production was proportional to brain mass over the entire lifespan, and therefore highest per body mass among young children^66^. Among prepubertal children (7 - 9 years of age), the rate of gluconeogenesis was still 50% higher than that found among adolescents^67^. The evolutionary plasticity of the liver appears to have co-opted the liver’s inherent developmental plasticity: among mammalian organs, only the liver retains full regenerative potential^25^. If the evolution of liver size has been determined by the demand for glucose production, then situations of high energy reliability/low dependence on gluconeogenesis in incipient clades may have favored evolutionary decreases in liver size, such as those associated with the divergence of Lagomorpha from Glires, and of Catarrhini from Haplorhini. Conversely, smaller bodies and tradeoffs favoring the liver appear to signify adaptation to limited energy resources relative to population size, exemplified by the divergence of Glires from Euarchontoglires, Strepsirrhini from primates, Platyrrhini from Haplorhini, and Callitrichinae from Platyrrhini.

Hippo-YAP, the regulator of organ size in Bilateria^68^, coordinates liver growth with the suppression of gluconeogenesis^69-72^. At low cell densities, Hippo signaling is normally suppressed, allowing YAP to localize in the nucleus and stimulate cellular proliferation, while at normal organ size and tissue density, YAP translocation and activity are normally inhibited by Hippo^68^. The global underexpression of microRNAs that is frequently observed in tumor cells has been attributed to aberrant density-dependent regulation of the same Hippo-YAP signaling: constitutively active YAP suppresses microRNA levels in cells and tumors and post-transcriptionally induces oncogenic MYC protein^73-74^. By extension, the evolutionary plasticity of the liver implies that each species divergence involves a reset of the dynamic ranges for the interactions among Hippo-YAP signaling, organ size, and the rate of microRNA biogenesis.

### A feedback model for brain size and energy reliability

We modeled a simple feedback loop based on the inverse association between larger brain sizes and *mirFam* and lower liver sizes and *cMR*s, assuming that the latter reflects decreased dependence on gluconeogenesis: a larger brain size leads to greater energy reliability which, in turn, facilitates the evolution of a larger brain, up to the limits imposed by anatomy and physiology, such as the diameter of the birth canal or the physiological limit on the rate of heat dissipation^10-12^. We applied a logistic feedback function to fit the distribution of *mirFam*, a proxy for energy reliability, to the primate brain size dataset of Grabowski et al (2023)^50^:

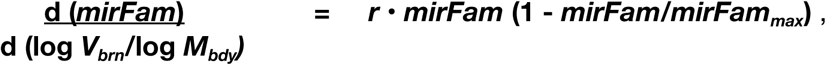

where ***r*** is the growth rate, and the range of *mirFam* has been scaled to a range of 1.0.

As shown in Figure 10-A, log *Mbrn*/log *Mbdy* is a predictor of *mirFam,* rather than the converse, consistent with the notion that evolutionary increases in brain size tended to result in greater energy reliability. Since the distribution of *mirFam* can be fit to the mass log ratios of any thermogenic organ, we asked which relationship might be the principal target of selection. Since the ranges of *mirFam* are highly clade-specific (Supplementary Figure S5) it is not a suitable dependent variable for phylogenetic regression. Instead, we inverted the premise and analyzed the dependence of organ *cMR*s on *mirFam*, as the independent variable.

**Figure 10.**
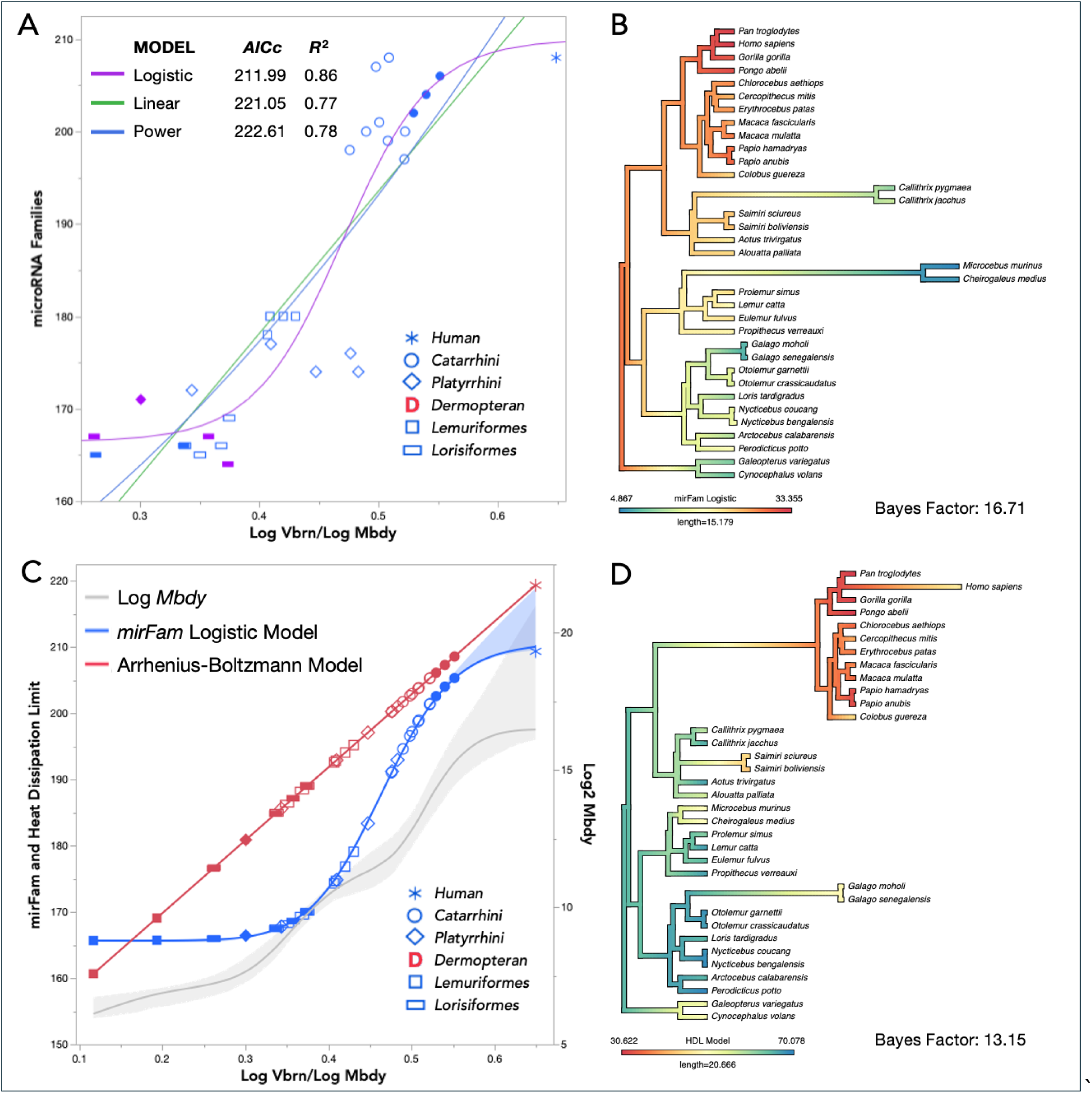
Logistic model of the *mirFam* relationship to relative brain size and the primate heat dissipation limit. **A.** Comparison of the logistic model with linear and power law models (*N* = 31, excluding the clade of heterothermic Cheirogaleidae: *M. murinus* and *C. medius*). Filled symbols: circles = Hominidae; rectangles and diamond: documented (blue) and suspected (purple) heterotherms. **B.** Variable rate phenogram based on the linearized values of the logistic model: when ranked in ascending order, the ratio of *mirFam* values to the cumulative sum decreases in a linear manner (Hubbert, 1982)^75^. **C.** Comparison of the logistic models with the maximal rate of thermogenesis predicted from whole-body mass and *T*_b_ (Allen et al, 2006)^76^ and scaled to the same range as *mirFam*. Shading represents the CI (cubic spline method). **D.** Variable rate phenogram based on the difference between predicted *mirFam* and model HDLs. The resulting distribution was centered and the outlier clade of dwarf and mouse lemurs was assigned an intermediate value of 50.

All potential kidney and heart *cMR*s (*cMR.60* - *cMR.80*) varied independently of phylogenetic distance, as did all but one liver *cMR* (*cMR.60lvr*) (Supplementary Table S5). In contrast, brain *cMR* variation with respect to *mirFam* was, in every case, secondary to phylogenetic distance (Pagel’s *lambda* = 1.0); in other words, the divergence of primate clades coincided with changes in *mirFam* range specifically with respect to brain *cMR*.

A variable rate phenogram corresponding to this model was generated using ratios from the linearization of the logistic function^75^, such that the color scale signifies rank order (Figure 10-B). Catarrhini exhibited the highest *mirFam* levels with respect to brain size, a relationship that evolved at the same rate as the underlying rate of genomic evolution. In contrast, the three smallest-bodied clades, *Cheirogaleidae, Galago and Callithrix* (marmosets) exhibited lower *mirFam* levels and an accelerated descent in rank (Figure 10-B). The latter observation is consistent with reversion of the brain size allometry of Callitrichines to one more similar to that of Strepsirrhini than that of Platyrrhini^64^. *Callithrix pygmaea* may also be heterothermic^61^, and the *Callithrix* clade also exhibited marked increases in the level and rate of evolution of liver *cMR.80*, suggesting adaptation to a regime of reduced energy reliability that is characteristic of low calorie-density folivory or insectivory^77^.

Supported by the previous observation that larger-bodied primates and Carnivora occupy the lower end of the range of Core/body ratios, we postulated that the upper plateau of the logistic model would correspond to the physiological limit on the rate of heat dissipation^10-12^. Furthermore, when primate species are ranked by size, *mirFam* varies reciprocally with inverse body temperature, 1/*T*_b_ (°*K*); thus, the seemingly random variation of *T*_b_ within each clade appears to be tied to the variation in *mirFam*, except among the smallest heterotherms (Supplementary Figure S6). The conserved nature of this reciprocity lends itself to the following formulation of a model of the maximum heat dissipation limit (HDL). The relationship between whole body *rMR* and *T*_b_ can be described by the Arrhenius-Boltzmann function, e^-*E*/*kT*^, where *E* represents the mean activation energy of the respiratory complexes and *k* is the Boltzmann constant^76^; when corrected for total body mass, this function represents the maximal possible rate of thermogenesis and therefore heat dissipation, regardless of phylogeny or thermoregulatory adaptations (Figure 10-C).

In order to detect adaptations that occurred in relation to this limit, we generated variable rate phenograms that depict the direction and rate of evolution of the *difference* between the values of the HDL model and the logistic model (Figure 10-C). In this case, the color scale signifies proximity to the HDL. Interestingly, two of the clades that exhibit accelerated evolution toward the HDL represent the highest relative brain size among Platyrrhini (*Saimiri*) and lowest relative brain size among Lorisiformes (*Galago*), suggesting thermoregulatory adaptation at the extremes of variation (Figure 10-D) (Supplementary Table S2). The divergence of Catarrhini was associated with rapid evolution toward the HDL that was subsequently reversed in two cases: the divergence of the human and colobus monkey, species that again represent the extremes of relative brain size within their clade (Figure 10-D). The apparent “cooling” in the human branch appears to reconcile the disproportionate increase in brain size with our species’ unique adaptations for heat dissipation^18^.

Our logistic model assigns the ancestral primate a high rank for energy reliability and a low rank relative to the HDL, a propitious combination for the founder of a new order of mammals. In many respects this model fulfills the recent call for an explanation of the distinct pattern of brain size evolution in primates “in terms of the release of a constraint or an adaptive shift that initiated escalating feedback between brain and behavior”^4^, assuming that such behavior results in greater energy reliability, hence reproductive fitness. Since this logistic model is agnostic regarding underlying mechanisms, it can also serve as a null hypothesis for comparison with the evolution of other candidate mechanisms of brain size evolution. More significantly, discovery of the tradeoff proposed four decades ago in order to account for thermal homeostasis during the evolutionary expansion of brain size^22-23^ depended on the definition of variables that are more sensitive to thermoregulatory constraints: arithmetic ratios, rather than log ratios or log-based slopes, and cellular *MR*s matched to an index of energy turnover and independent of developmental control, rather than log-based specific *MR*s.

## SYNTHESIS

The adaptive value of repression by individual microRNAs operating in gene regulatory networks has been attributed to their ability to canalize phenotype expression by buffering the effects of environmental or allelic variation^78-79^ and to set thresholds for cell fate determination, or maintain the identity of differentiated cells, by defining ranges of protein expression^80-81^. Major stages in the diversification of animal cell types, morphologies, and complexity of behavior have corresponded to significant expansions of microRNA and target repertoires^82-86^. For example, the emergence of placental mammals is marked by the appearance of 13 novel implantation-related microRNA families^33^, and the cognitive sophistication of the genus *Octopus* has been linked to 34 novel microRNA families that are preferentially expressed in neural tissues^82^. However, these *post hoc* associations do not shed light on the preconditions of repertoire expansion, nor on how such enhanced traits may be related to a higher range of *mirFam*, as opposed to action of individual novel microRNAs^82^. The consistent pairing of adaptive shifts in *MR*/kg with expansions of the microRNA repertoire implies a prerequisite of a high level of energy reliability in an incipient clade. Given that *mirFam* losses have occurred much less frequently than gains^31^, these observations suggest the operation of a ratchet mechanism that selects, among novel microRNAs, those that confer greater energy reliability. An analogy may be drawn to the fate of the duplicated genes of transcription factors (TF): owing to their established specificities, they have been much more likely to be retained than the paralogues of non-TF genes^87^.

## METHODS

### Physiological Data

The masses of the four thermogenic organs were obtained from the 111-species dataset of Wang et al (2012)^50^, originally derived from the compilation of Navarette et al (2011)^24^. Only in humans have the metabolic rates of the four thermogenic organs been measured directly^51^. Otherwise, they have been inferred by conventional allometric scaling based on organ mass of rat (all four organs), dog (liver, brain and kidney), cat and rabbit (liver)^50, 52^. Relative brain sizes for the logistic models were obtained from Grabowski et al (2023)^54^, Smaers et al (2021) (human)^64^, and estimated for Dermoptera^88^.

Values for thermal conductance (*C*_min_) were obtained from the dataset of Fristoe et al (2015)^6^, with substitution of congeneric species from the dataset of Wang et al (2012)^50^. Where *C*_min_ was available for two or more congeneric species, we chose the one with the mass nearest to the species in the Wang dataset. Body temperatures were obtained from the AnAge database^89^ and other sources.

### MicroRNA Families, Binning

Complements of *conserved* microRNA families were obtained from the most complete assembly for each species using the covariance model MirMachine^90^, trained on the manually-curated database of microRNA genes, MirGeneDB2.1^26^ (Supplementary Tables S1 and S2). For Scandentia, Lagamorpha and Rodentia, *mirFam* refers to the number of families common to Euarchontoglires. For Lorisiformes, Lemuriformes and Platyrrhini, *mirFam* refers to the number of families shared with Catarrhini. For the original regression of organ *cMR*s on *mirFam* and organ size (Figure 3-A), *mirFam* was binned according to natural divisions in its distribution: 141 - 161 (Scandentia and Glires), 165 - 181 (Glires, Lemuriformes and Platyrrhini), and 194 – 208 ‘(Catarrhini) (Supplementary Figure S5).

### Phylogenetic Trees and Variable Rate Phenograms

Using Mesquite © (2021) we derived ultrametric trees for 108 Theria, 51 Euarchontoglires or 24 primates from the mammalian phylogeny of Alvarez-Carretero et al (2021)^48^. Variable rate trees and ancestral node values were inferred using BayesTraits (version 4)^91-92^ (software available at www.evolution.rdg.ac.uk). Phenograms were generated using Phytools^93^.

For variable rate trees, Bayes factors were computed from the difference between the median of the top 1% likelihoods and the median likelihood of the 1% of trees with the smallest non-zero sum of changes to branch length, *i.e.* those trees most similar to the reference time tree, as previously described^12^. In the case of those variable rate trees based on linearization of the logistic function^75^, artifactual changes of slope at the extremes were avoided by the addition of dummy values to either end of the range.

### Statistics

Linear, logistic, and partial least squares regressions^57-58^ and normal mixture clustering^59^ were carried out and visualized using JMP16®. All *R*^2^ values were adjusted for the number of predictors. Other than linear and logistic regressions, curves were fit empirically using the cubic spline method, with the confidence intervals indicated by shading^94-95^. Phylogenetic linear regressions were carried out using the R: CAPER package^96^.

### Limitations

The three principal limitations of this study are:

i. the indirect origin of the mammalian organ *rMR*s, as described above (*Physiological Data*)^50, 52^.
ii. the small number and variable quality of the body temperature data in the AnAge database^89^;
iii. underestimation of the number of conserved microRNA families by the MirMachine algorithm^90^, relative to the reference, manually-curated database^26-27^ (currently MirGeneDB 3.0). Excluding Chiroptera, the correlation of computed values with MirGeneDB 3.0 is between 90 - 100% for genomes with scaffold N50 > 10,000^90^.

None of these factors is expected to affect the qualitative interpretation of our results.

## Supporting information

SUPPLEMENT Sorger and Fromm 2025

## DECLARATIONS

### Acknowledgment

BF is supported by the Tromsø forskningsstiftelse (TFS) [20_SG_BF ‘MIRevolution’].

### Author Contributions

Conceptualization: TS; Methodology: TS; Investigation: BF and TS; Writing – Original Draft: TS; Writing – Review & Editing: BF and TS; Visualization: TS; Funding Acquisition: BF.

### Statement on the use of AI

No part of this work was generated or edited using any Large Language Module.

