## SUPPLEMENT Sorger and Fromm 2025 for "Clockwork Orangutan: microRNAs, thermoregulatory tradeoffs, and models of brain size evolution"

**<u>SUPPLEMENTARY INFORMATION</u>**

Thomas Sorger, PhD*
Department of Biology
Roger Williams University
Bristol, RI 02809

Bastian Fromm, PhD**
UiT - The Arctic University of Norway,
The Arctic University Museum of Norway
9006 Tromsø, NO

* Lead contact
** Second corresponding author

**SUPPLEMENTARY INFORMATION**

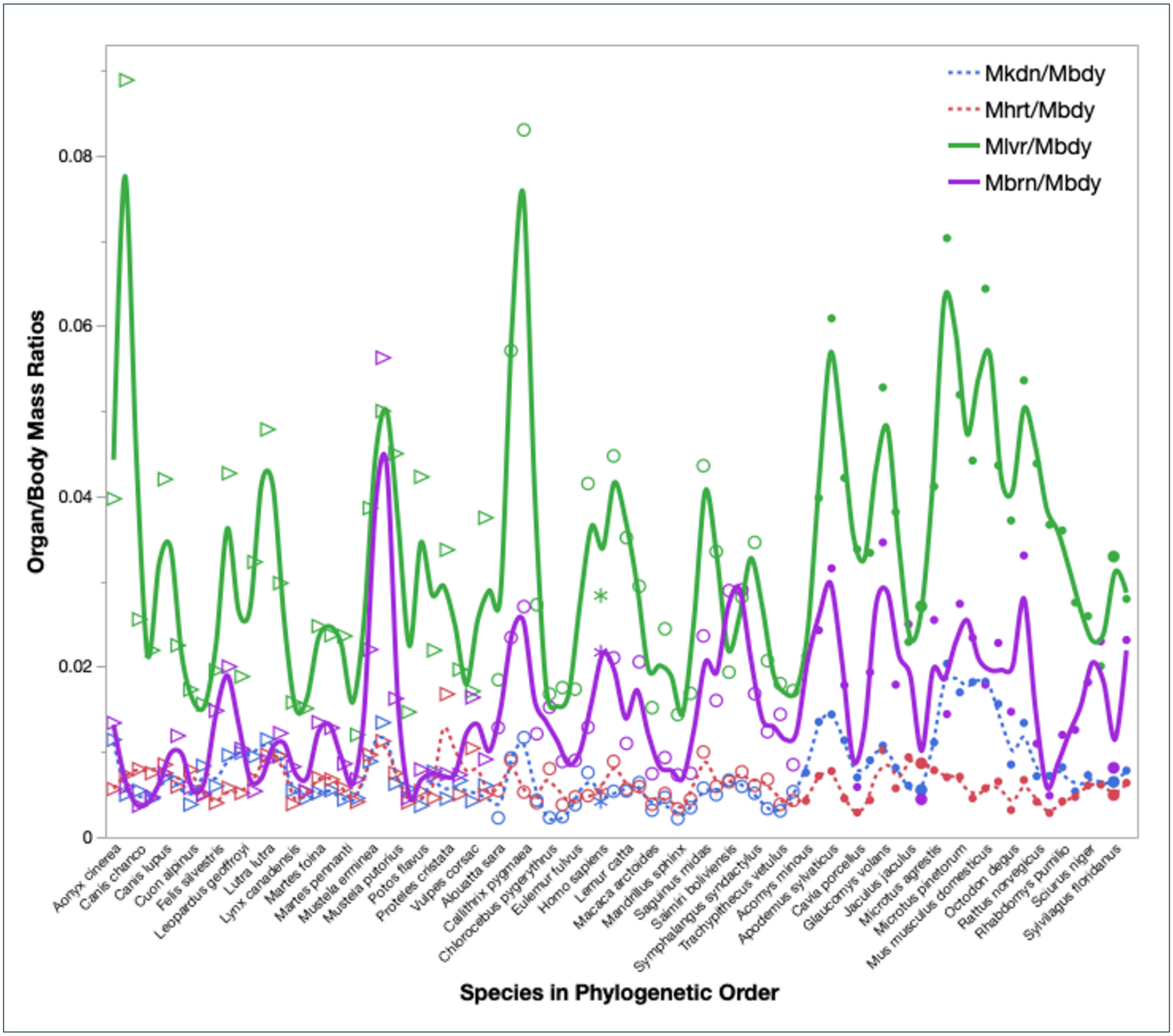


**Figure S1**. Organ/body mass ratios of the four thermogenic organs covary independently of phylogeny. Data point represent 80 species in the orders Carnivora, Primata and Glires, of which 40 have been labelled.

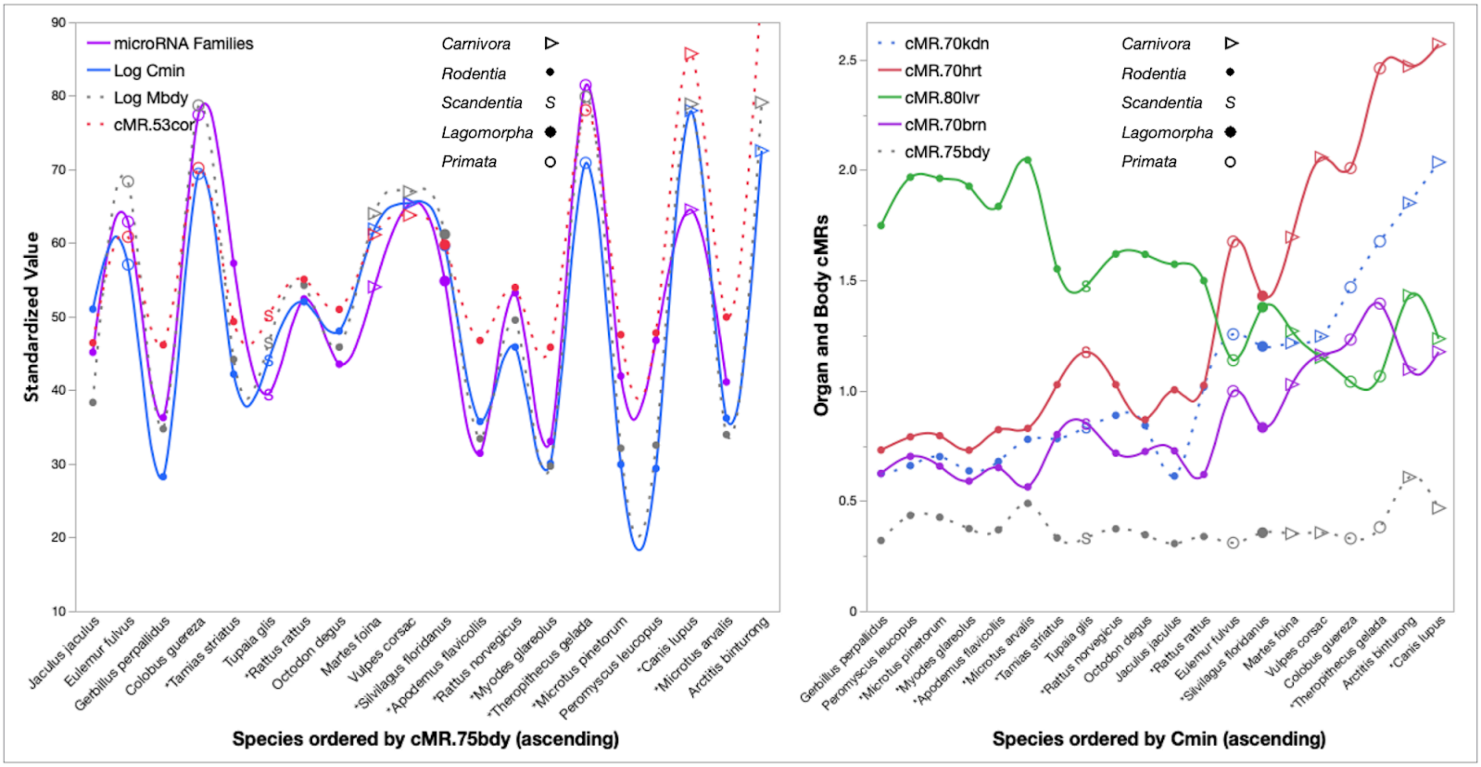


**Figure S2**. Thermoregulatory relationships among thermal conductance ($C_{min}$) , *mirFam* and organ *cMR*s.
A. *mirFam* tracks $C_{min}$ when mammals are ordered according whole body *cMR.75*. *cMR.53cor* represents the predicted rate of thermogenesis of the thermoregulatory Core (see main text Figure 6-A).
B. When ordered by thermal conductance ($C_{min}$), tradeoffs are apparent between liver *cMR.80*, on the one hand, and brain and heart *cMR.70*, on the other. *Asterisks indicate species in the 4-organ dataset of Wang et al (2012)[2] that have been substituted for congeneric species in the $C_{min}$ dataset of Fristoe et al (2015)[3].

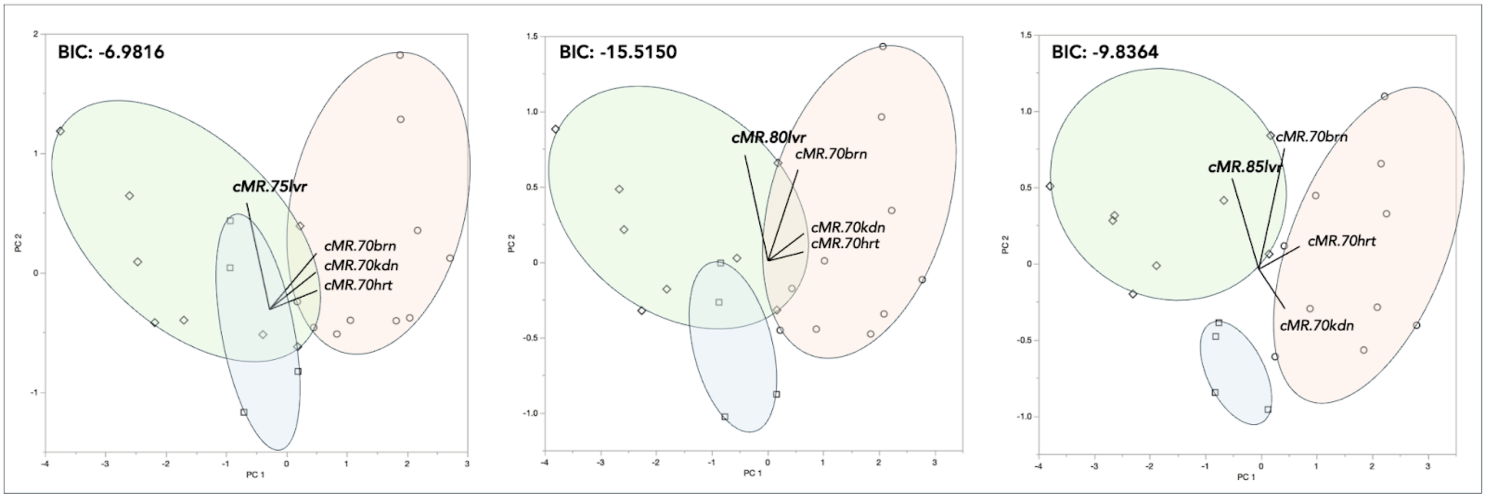


**Figure S3**. Unsupervised classification of primate clades based on Core *cMR* composition.
Of the *cMR* models with *VIP* = 4 (Figure 6-B), only two (with *cMR.80lvr* and *cMR.85lvr*) resolved Platyrrhini (diamonds) from Lemuriformes (squares) and Catarrhini (circles). Liver *cMR* varied between *cMR.75lvr*-*cMR.85lvr* (***bold type***).
BIC: Bayes Information Criterion. Ellipses were hand-drawn to include all members of each clade.
Principal components were calculated from each correlation matrix and species were classified by expectation maximization of their joint probability distribution (normal mixtures)[4].

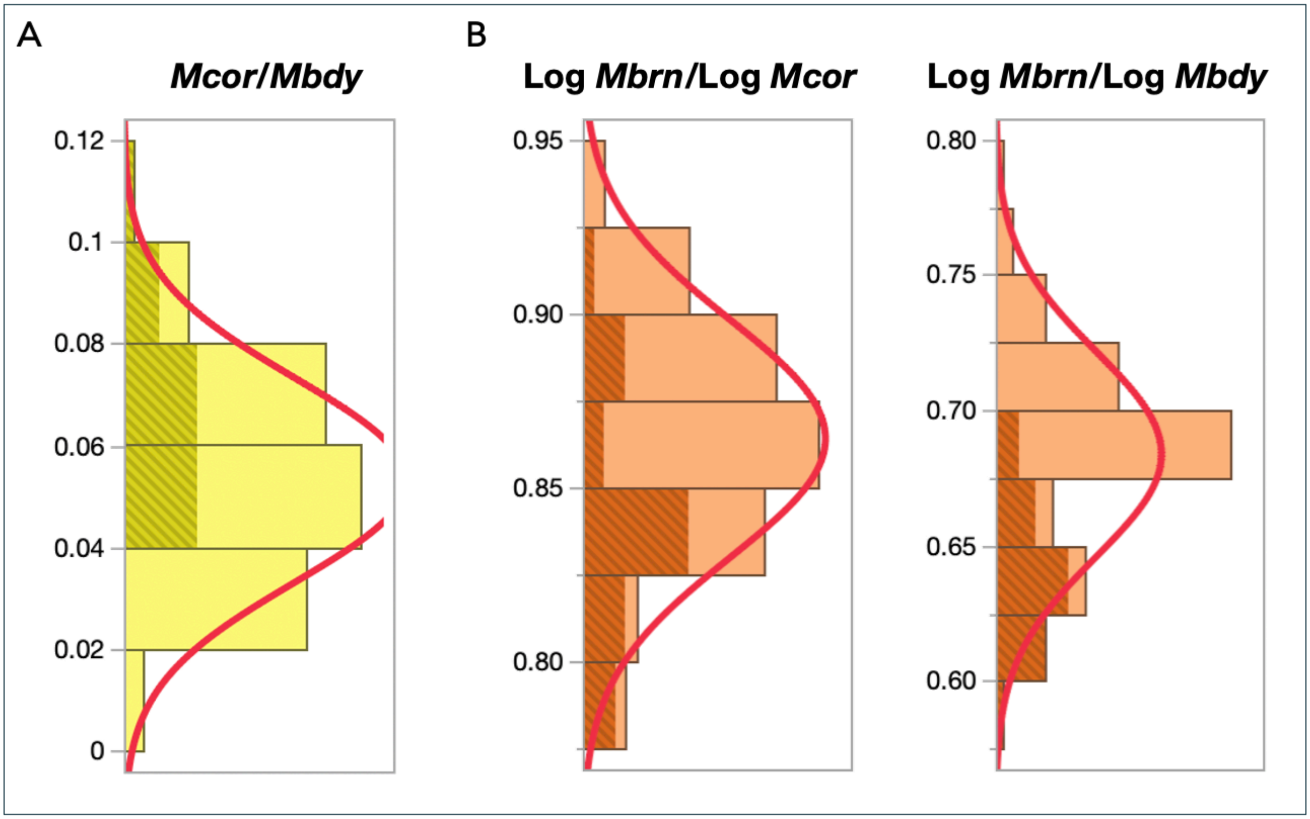


**Figure S4**. Distributions of arithmetic and log ratios among brain, core and whole body.

A. *Mcor/Mbdy* ratios in Boreoeutheria conform to a normal distribution (curve).

Note that Glires (grey) occupy the top of the range, consistent with an adaptation to maintain body temperature, while Primata and Carnivora occupy the lower end of the range, consistent with a heat dissipation constraint.

*N* = 78, excluding four outliers in upper 10 percentile of the range of values.

B. Log ratios with respect to Core mass yield a finer resolution of differences than log ratios based on body mass.

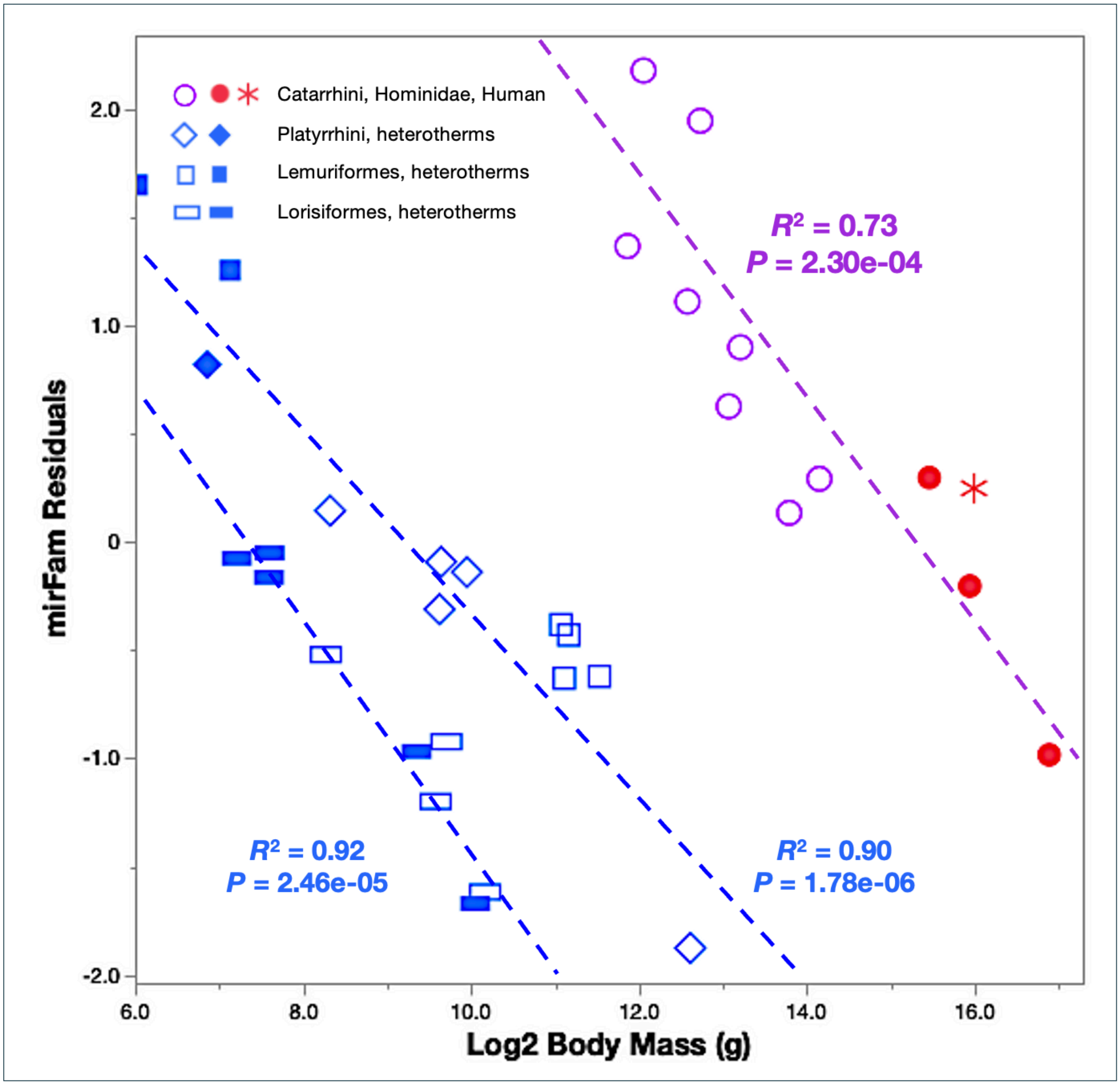


**Figure S5**. The ranges of *mirFam* are highly clade-specific and least conserved among Catarrhini.
For the regressions, species were grouped according to the natural divisions in the distribution of *mirFam*: Lorisiformes: 165 - 170; Lemuriformes and Platyrrhini: 171 - 180; Catarrhini: 197 - 208.

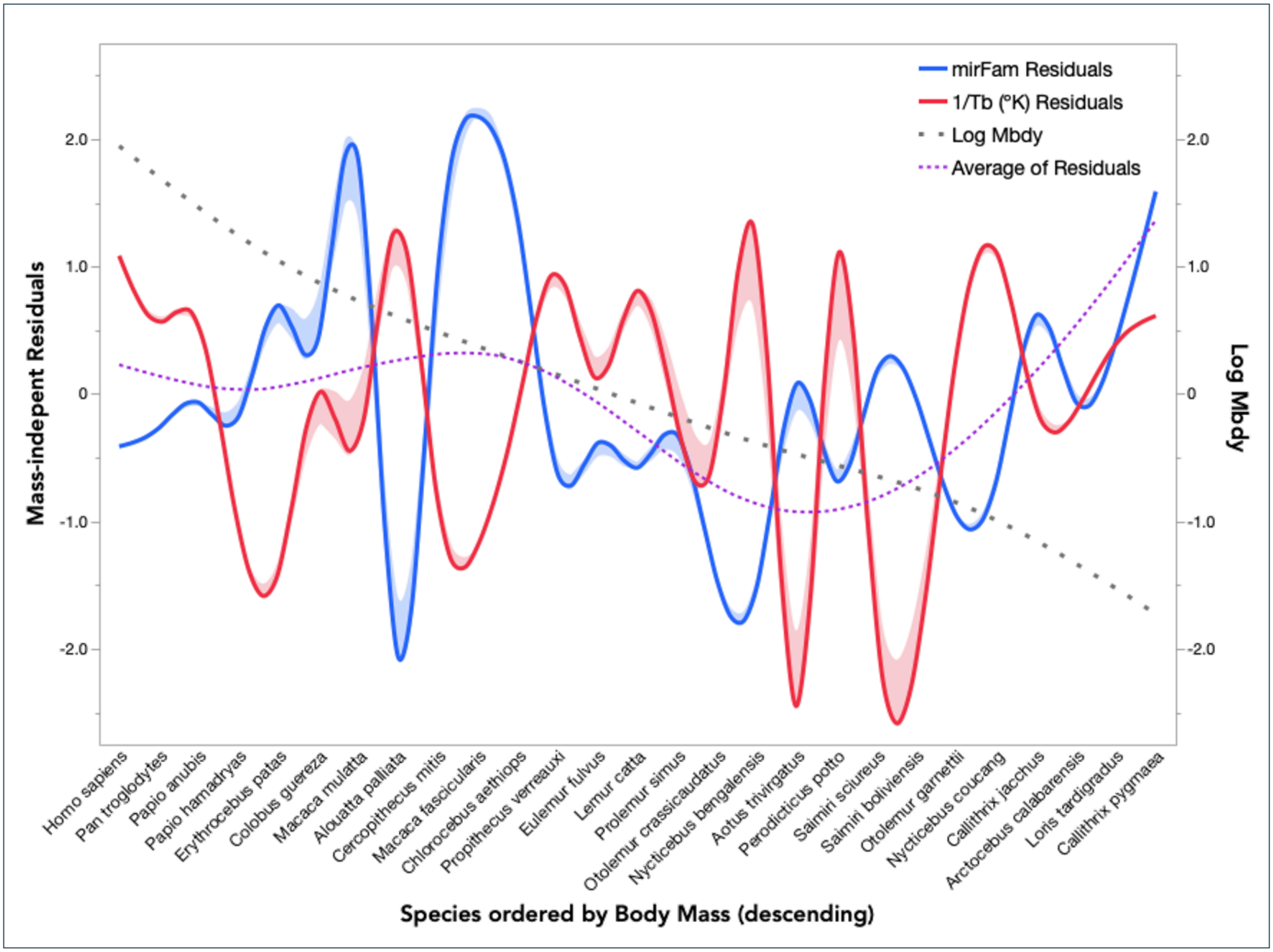


**Figure S6**. Reciprocal variation of *mirFam* and $1/T_b$ (°K) across primate clades (individual points omitted for clarity). Curves were fit empirically (cubic spline method) and the shading depicts confidence intervals (see Methods[5-6]). The original regressions on log *Mbdy* (g) excluded six outliers (gorilla, orangutan and four small heterotherms): ***mirFam***: $R^2$ = 0.70, $P$ = 4.01e-08, $N$ = 27; **$1/T_b$** (°K): $R^2$ = 0.36, $P$ = 9.87e-04, $N$ = 25 (two species lacked $T_b$ data).

**Supplementary Figure References**

1. Álvarez-Carretero S, Tamuri A, Battini M, Nascimento F, Carlisle E, Asher R, Yang Z, Donoghue P and Dos Reis M. 2022. A species-level timeline of mammal evolution integrating phylogenomic data. *Nature* **602**: 263-267.
2. Wang ZM, Zhang J, Ying Z and Heymsfield SB. 2012. Organ-tissue level model of resting energy expenditure across mammals: new insights into Kleiber's Law. *ISRN Zool*: 673050.
3. Fristoe TS, Burger JR, Balk MA, Khaliq I, Hof C and Brown JH. 2015. Metabolic heat production and thermal conductance are mass-independent adaptations to the thermal environment in birds and mammals. *Proc Natl Acad Sci USA* **112**: 15934-15939.
4. Celeux G and Govaert G. 1995. Gaussian Parsimonious Clustering Models. *Pattern Recognition* **28**: 781-793.
5. Reinsch CH. 1967. Smoothing by Spline Functions. Numerische Mathematik **10**: 177–183.
6. Eubank RL. 1999. Nonparametric regression and spline smoothing (2nd ed). CRC. Boca Raton, Florida.

**Table S1**. microRNA families, brain volume, body mass and temperature.
Endocranial volume (ECV) and body mass were obtained from Grabowski et al (2023)[1], Smaers et al (2021)[2] (human), or estimated from Martin (1995)[3] (Dermoptera).
Unless otherwise indicated, temperatures were retrieved from the AnAge database (Magalhães et al, 2024)[4].

| Order | Species | *mirFam* | ECV | Mass (g) | $T_b$ (ºC) | $T_b$ Source |
|---|---|---|---|---|---|---|
| Catarrhini | *Pan troglodytes* | 206 | 368.35 | 44967 | 37.2 | Jensen et al (2009)[5] |
| Catarrhini | *Homo sapiens* | 208 | 1252.00 | 50000 | 37.0 | |
| Catarrhini | *Gorilla gorilla* | 202 | 490.41 | 120950 | 35.5 | |
| Catarrhini | *Pongo abelii* | 204 | 65.00 | 3720 | 34.6 | |
| Catarrhini | *Chlorocebus aethiops* | 199 | 65.00 | 3720 | 36.8 | |
| Catarrhini | *Cercopithecus mitis* | 200 | 71.33 | 6109 | 37.5 | |
| Catarrhini | *Erythrocebus patas* | 201 | 97.73 | 9450 | 38.0 | Kolka and Elizondo (1983)[6] |
| Catarrhini | *Macaca fascicularis* | 207 | 63.98 | 4251 | 37.6 | |
| Catarrhini | *Macaca mulatta* | 208 | 88.98 | 6793 | 37.2 | Von Huben et al (2007)[7] |
| Catarrhini | *Papio hamadryas* | 197 | 146.17 | 14150 | 37.9 | Brain and Mitchell (1999)[8] |
| Catarrhini | *Papio anubis* | 200 | 167.42 | 18150 | 36.9 | |
| Catarrhini | *Colobus guereza* | 198 | 74.39 | 8589 | 37.0 | |
| Platyrrhini | *Callithrix pygmaea* | 171 | 4.17 | 116 | 35.2 | |
| Platyrrhini | *Callithrix jacchus* | 172 | 7.24 | 320 | 36.0 | |
| Platyrrhini | *Saimiri sciureus* | 176 | 24.14 | 799 | 37.5 | Robinson and Fuller (1999)[9] |
| Platyrrhini | *Saimiri boliviensis* | 174 | 25.07 | 789 | | |
| Platyrrhini | *Aotus trivirgatus* | 177 | 16.85 | 989 | 38.0 | |
| Platyrrhini | *Alouatta palliata* | 174 | 49.88 | 6250 | 36.0 | |
| Lemuriformes | *Microcebus murinus* | 174 | 1.63 | 65 | 37.3 | |
| Lemuriformes | *Cheirogaleus medius* | 176 | 2.60 | 140 | 38.0 | |
| Lemuriformes | *Prolemur simus* | 180 | 27.14 | 2150 | | |
| Lemuriformes | *Lemur catta* | 178 | 22.9 | 2210 | 36.0 | Larsen et al (2011)[10] |
| Lemuriformes | *Eulemur fulvus* | 180 | 25.77 | 2292 | 36.5 | |
| Lemuriformes | *Propithecus verreauxi* | 180 | 26.21 | 2955 | 36.0 | |
| Lorisiformes | *Galago moholi* | 165 | 3.71 | 148 | 38.0 | Nowack et al (2023)[11] |
| Lorisiformes | *Galago senegalensis* | 167 | 3.96 | 194 | 37.9 | |
| Lorisiformes | *Otolemur garnettii* | 166 | 11.50 | 764 | 36.0 | |
| Lorisiformes | *Otolemur crassicaudatus* | 165 | 11.78 | 1150 | 36.6 | |
| Lorisiformes | *Loris tardigradus* | 166 | 5.87 | 193 | 35.5 | |
| Lorisiformes | *Nycticebus coucang* | 167 | 10.13 | 653 | 35.4 | |
| Lorisiformes | *Nycticebus bengalensis* | 164 | 13.49 | 1060 | 35.5 | Loris Conservation Data[12] |
| Lorisiformes | *Arctocebus calabarensis* | 166 | 6.92 | 309 | 36.0 | |
| Lorisiformes | *Perodicticus potto* | 169 | 12.42 | 835 | 35.5 | Loris Conservation Data[12] |
| Dermoptera | *Galeopterus variegatus* | 177 | 6.9 | 1100 | | |
| Dermoptera | *Cynocephalus volans* | 173 | 6.9 | 1300 | | |

**Table S1 References**

1. Grabowski M, Kopperud BT, Tsuboi M and Hansen TF. 2023. Both diet and sociality affect primate brain-size evolution. *Syst Biol* **72**: 404-418.
2. Smaers JB, Rothman RS, Hudson DR . . (many others) . . Safi K. 2021. The evolution of mammalian brain size. *Sci Adv* **7**: eabe2101.
3. Martin RD. 1993. Primate origins: plugging the gaps. *Nature* 363: 223-234.

4. Magalhães JP, Abidi Z, dos Santos GA . . (many others) . . Ka Po To P. 2024. Human Ageing Genomic Resources: updates on key databases in ageing research. *Nucl Acids Res* **52**: D900-D908.
5. Jensen SA, Mundry R, Nunn CL, Boesch C and Leendertz FH. 2009. Non-invasive body temperature measurement of wild chimpanzees using fecal temperature decline. *J Wildlife Diseases* 45: 542-546.
6. Kolka MA and Elizondo RS. 1983. Thermoregulation in Erythrocebus patas: a thermal balance study. *J Appl Physiol Respir Environ Exerc Physiol* 55: 1603-1608.
7. Von Huben SN, Lay CC, Crean RD, Davis SA, Katner SN and Taffe MA. 2007. Impact of ambient temperature on hyperthermia induced by (±)3,4-methylenedioxymethamphetamine in Rhesus Macaques. *Neuropsychopharmacol* 32: 673-681.
8. Brain C and Mitchell D. 2009. Body temperature Changes in tree-ranging baboons (*Papio hamadryas ursinus*) in the Namib Desert, Namibia. *Intl J Primatol* 4: 585-598.
9. Robinson EL and Fuller CA.1999. Endogenous thermoregulatory rhythms of squirrel monkeys in thermoneutrality and cold. *Am J Physiol* 276: *R* 1397 – *R* 1407.
10. Larsen RS, Sauther ML and Cuozzo FP. 2011. Evaluation of modified techniques for immobilization of wild ring-tailed lemurs (*Lemur catta*). *J Zoo Wildlife Med* 42: 623-633.
11. Nowack J, Mzilikazi N and Dausmann KH. 2023. Saving energy *via* short and shallow torpor bouts. *J Therm Biol* 114: 103572.
12. loris-conservation.org

**Table S2**. MicroRNA families, body mass, body temperature and thermal conductance in mammals. In order to compare organ *MR*s, species marked (*) from the dataset of Wang et al (2012)[1] were substituted for congeneric species of similar mass in the $C_{min}$ dataset of Fristoe et al (2015)[2].

| Order | Species | *mirFam* | $C_{min}$ (ml $O_2$/h/ºC) | Mass (g) | $T_b$ (ºC) |
|---|---|---|---|---|---|
| Marsupialia | *Didelphis virginiae** | 141 | 83.97 | 2630 | 34.8 |
| Eulipotyphla | *Erinaceus europaeus** | 129 | 45.09 | 950 | 35.2 |
| Carnivora | *Arctictis biturong* | – | 125.34 | 10300 | 36.0 |
| Carnivora | *Martes foina* | 169 | 54.08 | 1410 | 37.0 |
| Carnivora | *Canis lupus** | 182 | 192.86 | 10000 | 36.0 |
| Carnivora | *Vulpes corsac* | 183 | 71.45 | 2080 | – |
| Rodentia | *Apodemus flavicollis** | 141 | 6.85 | 25.13 | 38.3 |
| Rodentia | *Myodes glareolus** | 143 | 4.37 | 15.36 | 37.6 |
| Rodentia | *Gerbillus perpallidus* | 147 | 3.79 | 29.98 | 38.8 |
| Rodentia | *Microtus arvalis** | 153 | 7.10 | 27.03 | 37.5 |
| Rodentia | *Microtus pinetorum** | 154 | 4.32 | 21.19 | 37.7 |
| Rodentia | *Octodon degus* | 156 | 18.08 | 129.21 | 37.6 |
| Rodentia | *Jaculus jaculus* | 158 | 22.90 | 48.04 | 37.0 |
| Rodentia | *Peromyscus leucopus* | 160 | 4.14 | 22.39 | 36.0 |
| Rodentia | *Rattus rattus** | 167 | 24.78 | 390 | 38.9 |
| Rodentia | *Rattus norvegicus** | 168 | 15.24 | 210 | 37.4 |
| Rodentia | *Tamias striatus** | 173 | 11.40 | 103.8 | 37.0 |
| Lagomorpha | *Sylvilagus floridanus** | 170 | 83.97 | 972 | 38.3 |
| Scandentia | *Tupaia glis* | 151 | 13.36 | 141.1 | 37.0 |
| Primata | *Eulemur fulvus* | 180 | 36.91 | 2500 | 36.5 |
| Primata | *Colobus guereza* | 198 | 97.88 | 9750 | 37.0 |
| Primata | *Theropithecus gelada** | 203 | 110 | 11400 | 37.0 |

**Table S2 References**

1. Wang ZM, Zhang J, Ying Z and Heymsfield SB. 2012. Organ-tissue level model of resting energy expenditure across mammals: new insights into Kleiber's Law. *ISRN Zool*: 673050.
2. Fristoe TS, Burger JR, Balk MA, Khaliq I, Hof C and Brown JH. 2015. Metabolic heat production and thermal conductance are mass-independent adaptations to the thermal environment in birds and mammals. *Proc Natl Acad Sci USA* **112**: 15934-15939.

**Table S3**. Derivation of Organ *rMR*s and mass from whole-body *rMR*s.
Relationships were inferred by applying the allometric models for 22 primates (Wang et al, 2012)[1] to the *bMR* data of 30 primates (Clarke et al, 2010)[2]. The models adopted to generate Figure 4 (main text) have been highlighted.

| INFERENCE | MODEL | | *AICc* | $R^2$ |
|---|---|---|---|---|
| *rMRcor* ~ *rMRbdy* (W) | Linear: | 0.7192 + 0.373 • *rMRbdy* | 41.88 | 0.9753 |
| " | Exponential: | 31.42 - 31.066 • $e^{-0.015 \cdot rMRbdy}$ | 40.72 | 0.9796 |
| *rMRlvr* ~ *rMRcor* (W) | Linear: | 0.2244 + 0.41108 • *rMRcor* | 5.29 | 0.9730 |
| " | Exponential: | 123.19 - 122.98 • $e^{-0.003414 \cdot rMRcor}$ | 8.28 | 0.9730 |
| *rMRbrn* ~ *rMRcor* (W) | Linear: | 0.0552 + 0.2048 • *rMRcor* | 6.33 | 0.8951 |
| " | Exponential: | 4.0829 - 4.23 • $e^{-0.0754 \cdot rMRcor}$ | 5.42 | 0.9123 |
| *Mlvr* (g) ~ *rMRlvr* (W) | Linear: | -44.7388 + 73.2747 • *rMRlvr* | 194.95 | 0.9729 |
| | Exponential: | -278.07 + 261.45 • $e^{0.17126 \cdot rMRlvr}$ | 175.15 | 0.9904 |
| *Mbrn* (g) ~ *rMRbrn* (W) | Linear: | -9.5065 + 71.2193 • *rMRbrn* | 151.33 | 0.9852 |
| | Exponential: | -159.39 + 158.1357 • $e^{0.30789 \cdot rMRbrn}$ | 133.05 | 0.9944 |

**Table S3 References**


1. Wang ZM, Zhang J, Ying Z and Heymsfield SB. 2012. Organ-tissue level model of resting energy expenditure across mammals: new insights into Kleiber's Law. *ISRN Zool*: 673050.
2. Clarke A, Rothery P and Isaac, NJB. 2010. Scaling of basal metabolic rate with body mass and temperature in mammals. J Animal Ecology 79: 610-619.


**Table S4**. Comparison of parameters for the Core *cMR* models of the major clades of Boreoeutheria.
Owing to a disproportionate reduction in body size, Glires exhibit the highest median and average ratios of *Mbrn/Mbdy*, but the lowest slope of log *Mbrn versus* log *Mbdy*. One outlier was omitted from each clade.

| Clade | N | *Mbdy* (g) (median) | Median *Mbrn/Mbdy* | Average *Mbrn/Mbdy* | Standard Deviation | Allometric Grade (slope) | Core *cMR* Exponent | Core Model $R^2$ |
|---|---|---|---|---|---|---|---|---|
| Carnivora | 29 | 5330 | 0.0083 | 0.0098 | 0.0048 | 0.7060 | 0.60 | 96.6015 |
| Glires | 25 | 50 | 0.0193 | 0.0196 | 0.0080 | 0.7272 | 0.40 | 99.0760 |
| Primates | 23 | 4200 | 0.0129 | 0.0154 | 0.0071 | 0.8000 | 0.53 | 98.0708 |

| | ***Core Model Organ Coefficients*** | | | | ***Core Model VIP Values*** | | | |
|---|---|---|---|---|---|---|---|---|
| **Clade** | ***Kdn*** | ***Hrt*** | ***Lvr*** | ***Brn*** | ***Kdn*** | ***Hrt*** | ***Lvr*** | ***Brn*** |
| Carnivora | 0.496 | 0.672 | 0.297 | 0.060 | 1.060 | 1.158 | 0.866 | 0.887 |
| Glires | 0.638 | 0.489 | 0.030 | -0.130 | 1.179 | 1.089 | 0.873 | 0.813 |
| Primata | 0.531 | 0.689 | 0.367 | 0.088 | 1.076 | 1.057 | 0.894 | 0.961 |

**Table S5**. Phylogenetic regressions of organ *cMR*s on species *mirFam* (N = 22).
The dataset excluded two outliers (the human and the heterotherm *Cheirogaleus medius*).

| | **Allometric Exponent** | **Pagel's *lambda*** | ***$R^2$*** | ***P*** | ***AICc*** |
|---|---|---|---|---|---|
| **KIDNEY** | 0.80 | 0 | 0.5366 | 6.39E-05 | -42.70 |
| | 0.75 | 0 | 0.5834 | 2.14E-05 | -20.09 |
| | 0.70 | 0 | 0.5958 | 1.56E-05 | -0.93 |
| | 0.65 | 0 | 0.5946 | 1.61E-05 | 16.22 |
| | 0.60 | 0 | 0.5859 | 2.00E-05 | 32.13 |
| | | **Pagel's *lambda*** | ***$R^2$*** | ***P*** | ***AICc*** |
| **HEART** | 0.80 | 0 | 0.4937 | 1.60E-04 | -38.02 |
| | 0.75 | 0 | 0.6476 | 3.83E-06 | -17.88 |
| | 0.70 | 0 | 0.6962 | 8.44E-07 | 0.63 |
| | 0.65 | 0 | 0.7133 | 4.69E-07 | 17.65 |
| | 0.60 | 0 | 0.7163 | 4.21E-07 | 33.66 |
| | | **Pagel's *lambda*** | ***$R^2$*** | ***P*** | ***AICc*** |
| **LIVER** | 0.80 | 0 | 0.5724 | 2.79E-05 | -15.22 |
| | 0.75 | 0 | 0.4011 | 9.28E-04 | -10.98 |
| | 0.70 | 0 | 0.0854 | 1.01E-01 | -2.86 |
| | 0.65 | 0 | -0.0344 | 5.89E-01 | 10.24 |
| | 0.60 | 0.962 | 0.0145 | 2.66E-01 | 27.34 |
| | | **Pagel's *lambda*** | ***$R^2$*** | ***P*** | ***AICc*** |
| **BRAIN** | 0.80 | 1 | 0.0329 | 2.05E-01 | -63.9 |
| | 0.75 | 1 | 0.107 | 7.55E-02 | -35.75 |
| | 0.70 | 1 | 0.1336 | 5.07E-02 | -10.58 |
| | 0.65 | 1 | 0.151 | 4.17E-02 | 11.76 |
| | 0.60 | 1 | 0.1596 | 3.71E-02 | 32.24 |

**Table S6**. Source data for Bayesian phylogenetic regressions (main text Table 1).
Data for endocranial volumes and body mass were drawn from Grabowski et al (2023)[1].
Resting *MR*s were obtained by applying the following model, derived from the same species in the dataset of Clarke et al (2010)[2]: $rMR = 93.9 - 93.37 \cdot e \wedge (-0.02 \cdot Mbdy\ (kg))$.

| Order | Species | *ECV* | *Mbdy* (g) | *ECV/Mbdy* | *rMR (W)* |
|---|---|---|---|---|---|
| Catarrhini | *Cercopithecus mitis* | 71.33 | 6,109 | 1.17 | 10.66 |
| Catarrhini | *Erythrocebus patas* | 97.73 | 9,450 | 1.03 | 14.98 |
| Catarrhini | *Papio ursinus* | 178 | 22,300 | 0.80 | 29.26 |
| Catarrhini | *Papio hamadryas* | 146.17 | 14,150 | 1.03 | 20.52 |
| Catarrhini | *Papio anubis* | 167.42 | 18,150 | 0.92 | 24.92 |
| Catarrhini | *Colobus guereza* | 74.39 | 8,589 | 0.87 | 13.90 |
| Platyrrhini | *Saguinus geoffroyi* | 10.14 | 517 | 1.96 | 1.55 |
| Platyrrhini | *Callithrix pygmaea* | 4.17 | 116 | 3.59 | 0.48 |
| Platyrrhini | *Callithrix jacchus* | 7.24 | 320 | 2.26 | 1.07 |
| Platyrrhini | *Saimiri sciureus* | 24.14 | 799 | 3.02 | 2.18 |
| Platyrrhini | *Aotus trivirgatus* | 16.85 | 989 | 1.70 | 2.58 |
| Lemuriformes | *Alouatta palliata* | 49.88 | 6,250 | 0.80 | 10.85 |
| Lemuriformes | *Cheirogaleus medius* | 2.60 | 140 | 1.86 | 0.56 |
| Lemuriformes | *Eulemur fulvus* | 25.77 | 2,292 | 1.12 | 4.96 |
| Lemuriformes | *Propithecus verreauxi* | 26.21 | 2,955 | 0.89 | 6.05 |
| Lorisiformes | *Galago senegalensis* | 3.96 | 194 | 2.04 | 0.72 |
| Lorisiformes | *Galago moholi* | 3.71 | 148 | 2.51 | 0.58 |
| Lorisiformes | *Euoticus elegantulus* | 5.53 | 274 | 2.02 | 0.95 |
| Lorisiformes | *Otolemur crassicaudatus* | 11.78 | 1,150 | 1.02 | 2.90 |
| Lorisiformes | *Otolemur garnettii* | 11.50 | 764 | 1.51 | 2.10 |
| Lorisiformes | *Galago demidoff* | 2.65 | 75 | 3.53 | 0.34 |
| Lorisiformes | *Nycticebus coucang* | 10.13 | 653 | 1.55 | 1.86 |
| Lorisiformes | *Loris tardigradus* | 5.87 | 193 | 3.04 | 0.72 |
| Lorisiformes | *Perodicticus potto* | 12.42 | 835 | 1.49 | 2.26 |
| Lorisiformes | *Arctocebus calabarensis* | 6.92 | 309 | 2.24 | 1.04 |

**Table S6 References**

1. Grabowski M, Kopperud BT, Tsuboi M and Hansen TF. 2023.
Both diet and sociality affect primate brain-size evolution. *Syst Biol* **72**: 404-418.

2. Clarke A, Rothery P and Isaac, NJB. 2010. Scaling of basal metabolic rate with body mass and temperature in mammals. J Animal Ecology 79: 610-619.
